# ER-anchored protein sorting controls the fate of two proteasome activators for intracellular organelle communication during proteotoxic stress

**DOI:** 10.1101/2023.12.11.571118

**Authors:** Gautier Langin, Margot Raffeiner, David Biermann, Mirita Franz-Wachtel, Daniela Spinti, Frederik Börnke, Boris Macek, Suayib Üstün

**Affiliations:** Faculty of Biology & Biotechnology, Ruhr-University Bochum, Bochum, Germany; Center for Plant Molecular Biology (ZMBP), University of Tübingen, Tübingen, Germany; Institute for Cell Biology, Department of Quantitative Proteomics, University of Tübingen, Tübingen, Germany Method section for MS/MS analysis; Leibniz-Institute of Vegetable and Ornamental Crops (IGZ), Großbeeren, Germany; Institute of Biochemistry and Biology, University of Potsdam, Potsdam, Germany

## Abstract

Proteotoxic stress, characterized by the accumulation of damaged proteins, poses a significant challenge to cellular homeostasis. To mitigate proteotoxicity eukaryotes employ the proteasome that is regulated by proteasome activators, e.g. transcription factors that promote gene expression of proteasome subunits. As proteotoxicity originates in different compartments, cells need to perceive signals from various locations. Understanding which components integrate signals to address proteotoxicity is essential to develop strategies to cope with proteotoxicity but remain elusive. Here, we identify that the proteasome autoregulatory feedback loop acts as a gatekeeper to facilitate the communication between nucleus and chloroplast. We reveal that the ER-anchored protein sorting system (ERAPS) controls the proteasomal degradation or nuclear translocation of proteasome activators NAC53 and NAC78. While both transcription factors activate the proteasome gene expression, they repress photosynthesis-associated nuclear genes during proteotoxicity through association with a conserved cis-element. Our data implicate a general trade-off between proteasome function and energy metabolism unravelling an unprecedented mechanism of how eukaryotic cells cope with proteotoxicity. Collectively, our discoveries provide a novel conceptual framework in which the proteasome autoregulatory feedback loop coordinates subcellular proteostasis and the trade-off between growth and defence.

## Main

Protein homeostasis, hereafter proteostasis, is defined as the synthesis of proteins and their regulated degradation. This intimate balance between protein synthesis and degradation is tightly regulated in all organisms to respond to several environmental stimuli^1,2^. Various perturbations can alter this equilibrium, leading to proteotoxic stress, the accumulation of aberrant proteins that cause ultimately irreversible cellular damage. These perturbations can range from ageing to pathological diseases such as Parkinson and Alzheimer as well as infection by pathogens^1,3^. The latter is a consequence of the onset and maintenance of defense responses during microbial infections and microbes’ ability to manipulate proteostasis to their own benefit^3,4^.

To mitigate deleterious effects due to excessive proteotoxic stress, cells employ various protein quality control machineries. One of the major protein quality control machineries across the tree of life is the ubiquitin-proteasome system (UPS). The UPS controls proteostasis through selective elimination of defective proteins and short-lived regulatory proteins^5^. To rapidly react to proteotoxic stress, all the constituents of the 26S proteasome are under a tight transcriptional control. All eukaryotic kingdoms possess transcription factors (TFs) required for proteasome gene activation^6^. Yeast utilizes Rpn4, mammals Nrf1/2 and plants the pair of NAC TFs (NAM, ATAF and CUC) NAC53/NAC78 as a conserved mechanism to cope with proteotoxicity. In yeast and animals, proteasome activators Rpn4 and Nrf1/2 have been shown to be UPS targets themselves, being degraded by the proteasome in the cytosol or ER^6^. Consequently, proteasome malfunction impairs their degradation leading to their stabilization and subsequent nuclear translocation to activate proteasome gene expression^6^. In plants, NAC78 mediates the transcriptional activation of proteasome promoters through the association with the proteasome regulatory cis element PRCE [TGGGC] ^7^. In addition, loss of the NAC53/78 TF pair rendered plants hypersensitive to proteasome inhibition^8^. Therefore, a similar proteasome regulatory feedback loop has been proposed in plants^6^; however, to date, no evidence supports proteasomal removal of NAC53/78.

The involvement of the proteasome in various cellular pathways^9–11^, suggests that the autoregulatory feedback loop is a central mechanism to precisely integrate and coordinate stress responses through proteotoxic stress sensing. Indeed, a growing body of evidence suggests that localized proteotoxic stress in organelles causes the activation of respective proteasome-mediated degradation pathways^12–16^. This implies a multi-layered signaling role of the proteasome in governing subcellular proteostasis. This aspect is particularly relevant for semi-autonomous organellar nuclear-encoded proteins, as their regulation also involves cytosolic proteasomal degradation prior to organellar import^17,18^. As such, proteasome activators could act as gatekeepers of the communication between the nucleus and energy-producing organelles to maintain subcellular proteostasis. This could link the proteasome directly to energy metabolism as well as to the trade-off between growth and defense. However, to date it is unknown whether proteasome activators exert these functions.

Here we report that proteasome activators, TFs NAC53 and 78, orchestrate the communication between the nucleus and chloroplast during various stress conditions. We reveal that ER-anchored protein sorting (ERAPS) controls the proteasomal degradation and nuclear translocation of NAC53 and NAC78. While both TFs activate the proteasome gene expression, they repress photosynthesis-associated nuclear genes during proteotoxicity. The trade-off between proteasome activation and photosynthesis downregulation seems to be a general feature as it occurs in response to various environmental and developmental cues.

### NAC53/78 are central integrators of various proteotoxic stress conditions

Proteotoxicity is triggered by a wide range of physicochemical stresses at different subcellular locations often leading to accumulation of proteins that are not imported into these organelles ^17,19^. To understand whether proteasome activators might play a role in integrating signals from different locations, we used the proteasome activator mutant *nac53-1 78-1*^8^ to disrupt cellular proteostasis through a variety of approaches (Fig.1a): (i) inhibiting the proteasome via bortezomib (BTZ) and bacterial infection with *Pseudomonas syringae pv. tomato DC3000* (Pst)^20,21^, (ii) inhibiting the segregase CDC48 (CB-5083)^16,22^, (iii) targeting HSP90 (GDA)^23^, which plays a critical role in the import and degradation of chloroplastic/mitochondrial precursors, (iv) inducing ER stress with DTT and TM, and (v) targeting semi-autonomous organelles such as chloroplasts and mitochondria (Lin, Cml, MV)^23^.

**Fig. 1.**
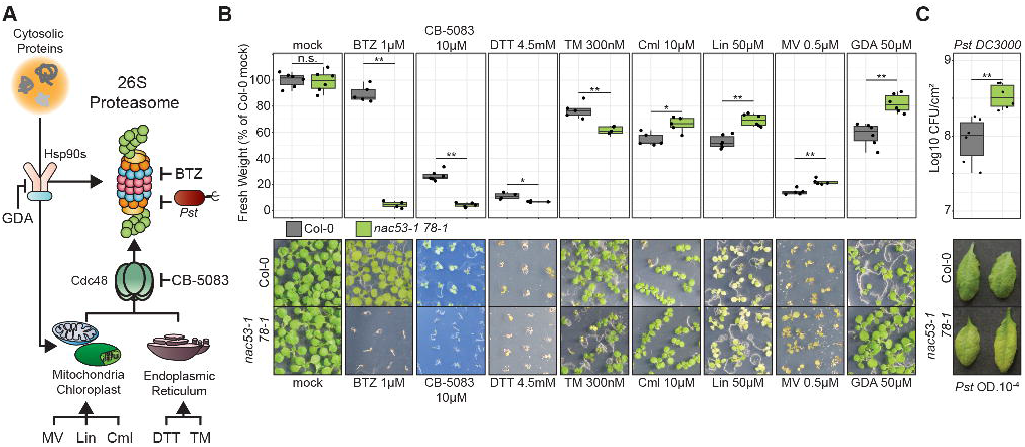
NAC53/78 are central integrators of various proteotoxic stress conditions. (A) The role of the 26S Proteasome in degradation of proteins from distinct subcellular compartments. Prime site of action of the several treatments used in this study: Blunt head arrows indicate enzymatic activity suppression; pointed head arrows indicate trigger of substrate accumulation. (B) Fresh weight of seedlings grown under the indicated treatments at 10-14 day after germination (dag). Fresh weight is expressed as a percentage of Col-0 mock conditions. Boxplots colors refer to the genotype. Statistical significance is assessed by a Wilcoxon-Mann-Whitney test (p values: n.s. > 0.05; * < 0.05; ** < 0.01). Every treatment has been repeated at least twice with similar results. (C) Bacterial density in Log10 Colony-Forming-Unit (CFU) per leaf cm². Statistical significance is assessed by a Wilcoxon-Mann-Whitney test (p values: n.s. > 0.05; ** < 0.01). Boxplots colors refer to the genotype as in panel B. The treatment has been repeated three times with consistent results. (B-C) Representative phenotypic pictures of Col-0 and *nac53-1 78-1* plants under several treatments are included.

Perturbation of the proteasome by chemicals, bacteria, CDC48 inhibition and induced ER stress resulted in an increased susceptibility of the *nac53-1 78-1* mutant (Fig.1b), suggesting that both proteasome activators are involved in a general regulation of subcellular proteostasis. Strikingly, all other treatments perturbing organelle function (Lin, Cml, MV, GDA) resulted in increased tolerance of *nac53-1 78-1* (Fig.1b). Overall, our phenotypic screen implies that NAC53 and NAC78 integrate signals from different compartments to adjust their impact on plant response to stress.

### NAC53/78 are part of an autoregulatory proteasome feedback loop

To elucidate how both TFs might integrate signals from different compartments and organelles, we aimed to decipher their regulation through interactome analysis. Transgenic Arabidopsis GFP-NAC53 and 78 lines (under the control of UBQ10 promoter) were only detectable for the GFP fusion proteins upon proteasome inhibition by BTZ (Fig. 2,b), implying a high proteasomal turnover of both NACs. Thus, we used *Agrobacterium*-mediated transient expression in *Nicotiana benthamiana* to boost the expression of NACs for our interactome studies. Transient expression of GFP-NAC53 and 78 (sTag construct) decorated the ER, being consistent with the predicted transmembrane domain (TM) at the C-terminus of both proteins (Fig. 2c and Extended Data Fig.1a). We additionally employed a chimeric construct (dTag-NACs) tagged with fluorophores at the N and C-terminus following the TM domain (Extended Data Fig.1a) to monitor subcellular interaction partners, as the RFP signal remained strictly in the ER (Fig. 2d). BTZ treatment, stabilized all NAC constructs, supporting the notion of NAC53/78 as proteasome targets and validating the functionality of the system in *N. benthamiana* (Extended Data Fig.1b,c).

Next, we performed immuno-precipitation (IP) followed by tandem mass spectrometry (MS/MS) with both constructs. We processed a total of 4 IP-MS/MS conditions for both NACs among two experiments. We performed IP of sTag-NACs in mock conditions and at 8 hours after bacterial infection, to induce proteotoxic stress (Extended data Fig.1D). In parallel, we performed IP of dTag-NACs enriching for the N- or C-terminus end of the proteins, which allowed us to characterize the interactome at a subcellular resolution (Extended data Fig.1E). For our downstream analysis, we focused on the common interactors for both NACs. Out of the 4 conditions we identified 245 unique *A. thaliana* orthologous proteins as interactor candidates (Fig. 2e, Supplementary Table 1). The Gene Ontology (GO) annotations for cellular components and the displayed enrichment for terms mainly associated with the proteasome complex and plastid components (Extended Data Fig.1f). Investigating the SUBA5 annotation of NAC53/78 interacting candidates, we found enrichment for following compartments: plastid, peroxisome, Golgi, and ER (Extended Data Fig.1g). Further GO analysis for biological processes revealed that possible candidates were enriched in proteins related to three main biological process: proteolysis, trafficking and translation (Fig. 2f). We extracted the corresponding proteins and performed a protein network analysis (Fig. 2g), highlighting the presence of UPS components (Fig. 2g; Extended Data Fig.1h-k). However, proteasome components were strongly reduced by bacterial infection and mainly enriched when targeting the ER-anchored end of the TFs (C-ter IP) (Extended Fig.1h), indicating that both proteasome regulators are degraded at the ER. The latter was further supported by the presence of several ER-associated degradation (ERAD) components in the interactome (Fig.2g and Extended Data Fig.1i). In summary, our interactome analysis validates the role of NAC53/78 as integrators of subcellular proteostasis disruptions, affirming their regulation by the proteasome autoregulatory feedback loop in plants.

**Fig. 2.**
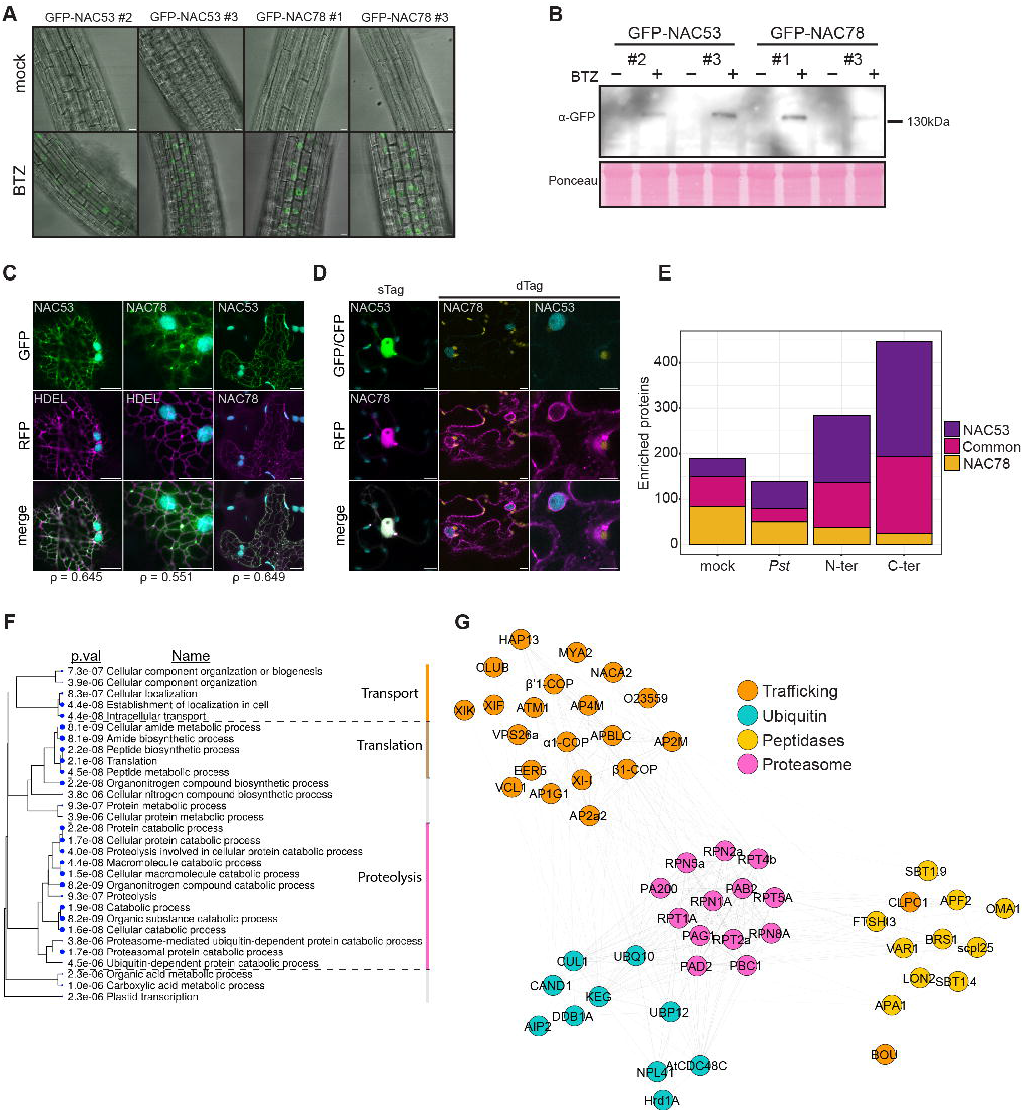
NAC53/78 are part of an autoregulatory proteasome feedback loop. (A) Confocal microscopy pictures of roots of GFP-NAC53/78 *Arabidopsis thaliana* transgenics at 7dag exposed to mock or BTZ 10µM treatment for 3h. Treatments were repeated at least three times with similar observations. (B) Immunoblot analysis against GFP on crude extract of seedlings related to panel A. Ponceau staining is used as loading control. (C) Confocal microscopy pictures of transiently co-expressed sTag-NAC53/78 with prom35S::RFP-HDEL or together in *N. benthamiana* epidermis cells. Pearson index (ρ) represents co-localization index. Scale bars = 10µm (D) Confocal microscopy pictures of transiently co-expressed sTag-NAC53/78 together or dTag-NAC53/78 alone in *N. benthamiana* epidermis cells. Scale bars = 10µm (E) Bar plots representing the number of significantly enriched proteins in the 4 IP-MS/MS conditions for NAC53 only, NAC78 only and NAC53+NAC78 (Common). (F) Cladogram of top 30 gene ontology (GO) terms for biological process (BP) enriched in the list of *A. thaliana* ortholog proteins found from NAC53/78 common interactome. (G) Protein network of the interacting candidates related to the GO BP terms annotated as “Transport” and “Proteolysis” in panel F.

### ER-anchored protein sorting (ERAPS) regulates the subcellular fate of NAC53 & NAC78

Our interactome approach revealed that NAC53/78 associated with candidates that belong to the ERAD machinery, including membrane bound E3 ligase HRD1 and CDC48 (Fig. 2g and Extended data Fig.1i). To test whether ERAD is required to turnover the NACs, we took advantage of pharmacological drugs targeting individual steps in ERAD: we treated the GFP-NAC53 and GFP-NAC78 transgenic lines with CB-5083 and LS-102, inhibitors of CDC48 and HRD1 proteins, respectively^22,24^. We confirmed that previously uncharacterized LS-102 can efficiently inhibit both *A. thaliana* orthologs AtHRD1a and AtHRD1b *in vitro* (Extended data Fig. 2a,b). Similar to BTZ and CDC48, HRD1 inhibition led to a stabilization of both proteins; confirmed by confocal microscopy and immunoblotting (Fig. 3a,b). Collectively, these results indicate that NAC53/78 are targets of ERAD. Interaction studies of NACs with HRD1 isoforms revealed that GFP-NACs strongly associated with RFP-HRD1a/b in localization and co-IP experiments (Fig. 3c,d). The association was largely reduced upon deletion of the TM domain of both NAC TFs (ΔTM, Fig. 3c). Interestingly, loss of ER association led to a strong stabilization of the proteins, supporting ERAD-dependent turnover for both TFs (Fig. 3e). Subsequent *in vitro* ubiquitination assays using purified GST-HRD1a/b isoforms and MBP-NAC53/78 proteins confirmed that both NACs are ubiquitinated by HRDs, indicated by the appearance of higher molecular weight bands of MBP-NAC proteins (Fig. 3f). We could confirm the presence of ubiquitination marks at different sites via MS/MS analysis of the *in vitro* ubiquitination reactions (Extended Data Fig. 2c,e). In addition, ubiquitin marks were consistently inhibited by the addition of the inhibitor LS-102 to the *in vitro* ubiquitination assay (Extended data Fig.2f). The *in vitro* data was corroborated by the identification of several ubiquitination sites on NAC53 and 78 using *in vivo* IP-MS/MS analysis (Extended data Fig. 2d,e). This could be confirmed by performing ubiquitin IP on *N. benthamiana* leaves transiently expressing the NAC constructs (Extended data Fig. 2g). Mutation of identified lysine residues substantially reduced the association with anti-ubiquitin beads *in planta* (**Extended data Fig. 2h,i**). Altogether, our *in vivo* and *in vitro* analysis demonstrate that NAC53 and NAC78 are directed to proteasomal degradation via ERAD through HRD1 ubiquitination.

**Fig. 3.**
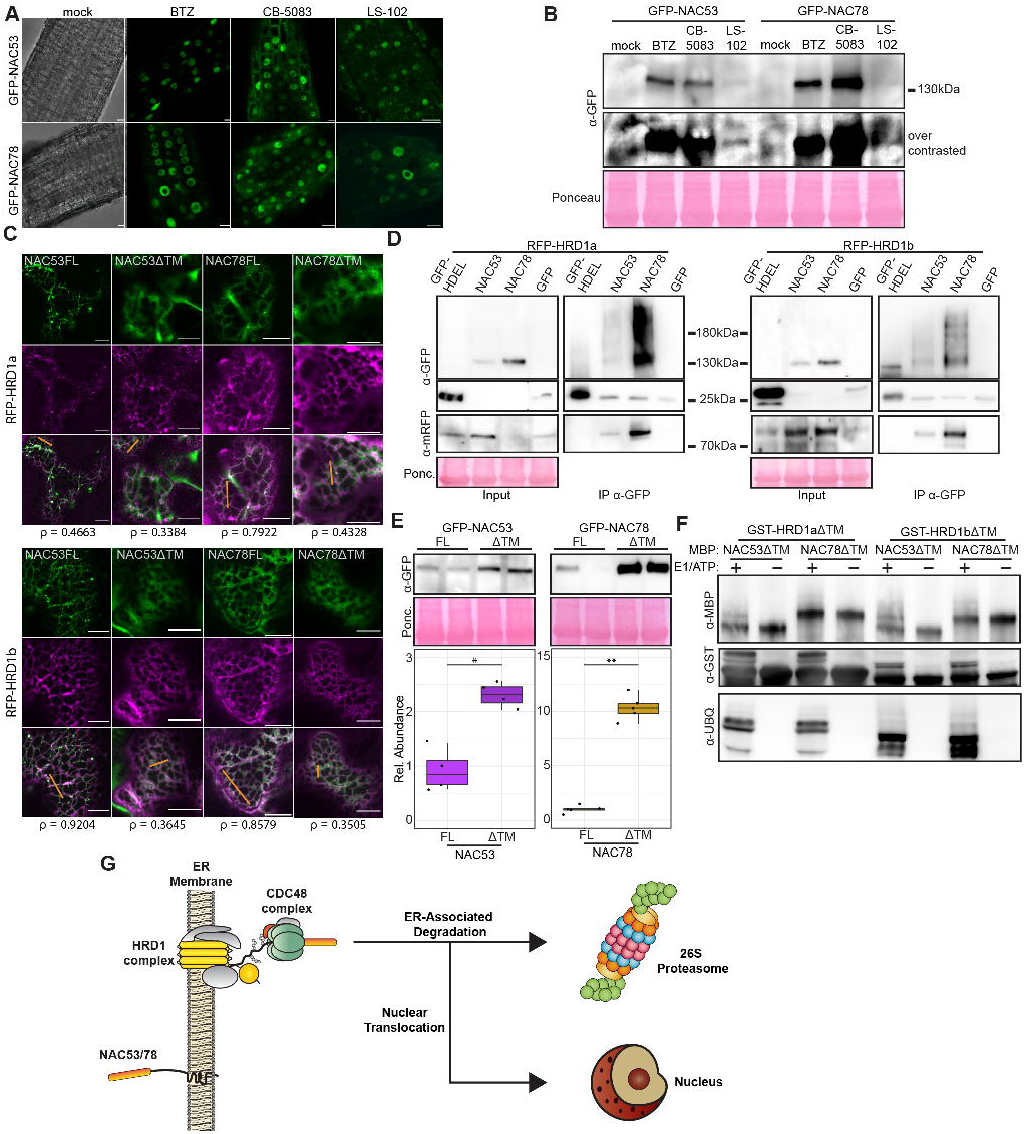
ER-Anchored Protein Sorting (ERAPS) coordinates the subcellular fate of NAC53 & NAC78. (A) Confocal microscopy pictures of GFP-NAC53/78 expressing *A. thaliana* roots. 7 dag seedlings were exposed to mock treatment, BTZ 10µM, CB-5083 10µM or LS-102 100µM for 3h. The treatments were repeated at least three times with similar observations. Scale bars = 10µm (B) Immunoblot against GFP on crude extracts of GFP-NAC53/78 *A. thaliana* adult plants infiltrated with mock treatment, BTZ 10µM, CB-5083 10µM or LS-102 100µM for 6h. Ponceau staining is used as loading control. (C) Confocal microscopy pictures of *N. benthamiana* transiently co-expressing GFP-NAC53/78 full-length (FL) or deleted for the transmembrane domain (ΔTM) with RFP-HRD1a or RFP-HRD1b. Pearson index (ρ) indicates correlation index at the orange lines. (D) Co-IP of RFP-HRD1a/b with GFP-NAC53/78. IP was performed with anti-GFP beads. GFP-HDEL and free-GFP were used as negative control. The experiment was repeated twice with similar results. (E) Immunoblot against GFP on crude extract of *N. benthamiana* leaves transiently expressing GFP-NAC53/78FL or ΔTM. Ponceau was used as loading control. Boxplots represent the relative ΔTM abundance compared FL. (F) *In vitro* trans-ubiquitination assays of GST-HRD1a/b against MBP-NAC53/78. Removal of E1 and ATP from the reaction was used as negative control. The experiment was repeated twice with similar results. (G) Representation of the ER-Anchored Protein Sorting (ERAPS) mechanism. NAC53/78 are ER-anchored proteins recognized and ubiquitinated by the HRD1 complex. The CDC48 complex subsequently extracts NAC53/78 from the ER to mediate ER-Associated Degradation or allowing nuclear translocation in the context of proteasomal stress to mitigate proteotoxicity.

During our analysis we observed that NAC53 and 78 responded different to the tested inhibitors: while BTZ led to a strong signal in the nucleus, CDC48 inhibition led to the appearance of cytoplasmic dot-like structures (Fig. 2a) that was not due to differences in drug potency depending on the concentration of the drugs (Extended data Fig. 3a). Co-expression of CDC48 isoform AtCDC48c with both NACs revealed that both proteins associated in dot-like structures (Extended data Fig. 3b), suggesting that NAC53/78 accumulate at sites of CDC48 action upon CDC48 inhibition. Given the role of CDC48 in retro-translocation of ER proteins to the cytosol we tested whether inhibition of its action can also perturb the subcellular sorting of NACs comparing the GFP signal intensity in the nucleus and cytoplasm upon BTZ or CB-5083 treatment. The proportion of the GFP signal in the nucleus was significantly lower when plants were treated with CB-5083, supported by the lowered ratio of nucleus/cytoplasm GFP intensity of plants subjected to the same treatment (Extended data Fig. 3c-f).

Together, our findings demonstrate the recycling of both NACs by the ERAD machinery, regulated by HRD1 and CDC48, with the latter facilitating their nuclear translocation (Fig. 3g). Thus, we propose to refer to this mechanism as ERAPS (ER-anchored protein sorting).

### The transcriptional landscape of proteotoxic stress

To understand why the *nac53-1 78-1* mutant is more tolerant to organelle-directed proteotoxic stress while it is hypersensitive to general proteotoxic stress we undertook a transcriptomic analysis. To this end we exploited the *Pst*-mediated proteotoxic stress caused by the suppression of proteasome activity and broad modification of chloroplast proteome^20,21,25^. Our analysis revealed that *Pst* induced proteotoxic stress hallmarks were dependent on NAC53/78 (Fig. 4a,b,c, Extended data Fig.4a) and stabilized both NAC TFs in *N. benthamiana* and *A. thaliana* (Fig. 4e,f). As such, this system is ideal to unveil the function of the proteasome autoregulatory feedback loop in response to environmental perturbation. Next, we analysed the transcriptomic profile of *nac53-1 78-1* infected with *Pst*, which revealed 945 differentially expressed genes (DEGs). Strikingly, apart from the proteasome cluster most of the DEGs displayed a positive Log2FC (Fig. 4g,h and Extended data Fig. 4a). This suggests that NAC53/78 are major transcriptional repressors upon bacteria-induced proteotoxic stress. Further investigation of which biological processes are enriched within the DEGs revealed photosynthesis, glucosinolates, cell wall and auxin signalling as major terms (Fig. 4i). To obtain a global view of the transcriptional landscape during proteotoxicity, we included two other datasets: (i) transcriptome analysis of proteasome mutant *rpt2a-2* in response to *Pseudomonas syringae pv. maculicola* (*Pma*) infection (Extended data Fig.4b-e) and (ii) previously characterized transcriptome analysis on Col-0 plants upon MG132 treatments^8^. The comparison of all three datasets allowed us to extract a core set of 538 DEGs present in the *nac53-1 78-1* dataset and at least one of the other datasets (Fig. 4j and Supplementary Table 2). Extracting the genes associated with these processes from the *nac53-1 78-1* 945 DEGs, we could define the core transcriptional network associated with proteotoxic stress (Fig. 4l). This network is comprised of 4 modules: the proteasome genes, a major cluster of photosynthesis associated nuclear genes (PhANGs), glucosinolate and auxin signalling pathways (Fig. 4k,l). Only the proteasome and photosynthesis cluster were present in all three datasets (Extended data Fig.4f). This suggests that PhANGs transcriptional repression is a recurrent response to proteotoxic stress. Altogether our analysis revealed a so far unidentified link between NAC53/78 and the transcriptional regulation of PhANGs in response to proteotoxic stress.

**Fig. 4.**
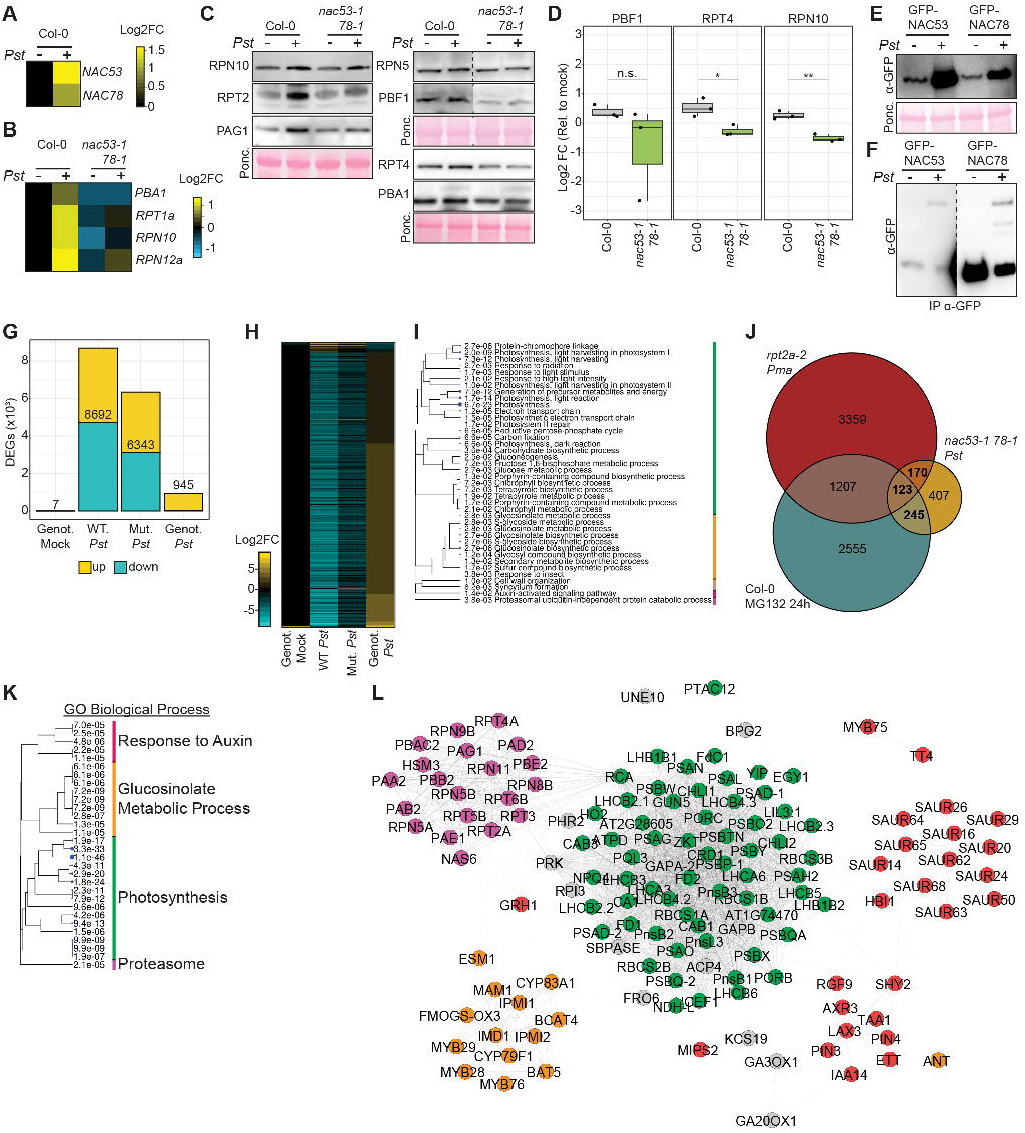
The transcriptional landscape of proteotoxic stress. (A) Log2FC mRNA levels of *NAC53* and *NAC78* transcripts after *Pst* infection of adult Col-0 plant leaves. The heatmap represent the mean of 4 biological replicates. (B) Log2FC mRNA levels of 26S proteasome transcripts after *Pst* infection of Col-0 or *nac53-1 78-1* adult plant leaves. The heatmap represent the mean of 4 biological replicates. The experiment has been repeated 3 times with consistent results. (C) Immunoblot against multiple 26S proteasome subunits on crude extracts of Col-0 or *nac53-1 78-1* adult plant leaves inoculated with *Pst* or mock solutions for 24h. Ponceau staining is used as loading control. (D) Relative Log2FC abundance of representative 26S proteasome subunits in Col-0 or *nac53-1 78-1* adult plant leaves. The abundance is calculated relative to the mock conditions. Statistical differences are assessed via a Welch t.test (p values: n.s > 0.05, * < 0.05, ** < 0.01). (E) Immunoblot against GFP on crude extract of *N. benthamiana* leaves transiently expressing GFP-NAC53/78 after mock treatment or *PstΔHopQ* infection for 8h. Ponceau staining is used as loading control. (F) Immunoblot against GFP on extract of adult *A. thaliana* transgenics GFP-NAC53/78 subjected to mock treatment or *Pst* infection 8h by vacuum infiltration followed by IP with anti-GFP beads. (G) Number of differentially expressed genes (DEGs) in the different conditions of the *nac53-1 78-1 Pst* RNAseq analysis. Total number of DEGs per condition is indicated. Colors indicate the proportion of DEGs up-regulated and down-regulated. DEGs are considered when |Log2FC| > 1.5. Genot. mock: *nac53-1 78-1* mock vs. Col-0 mock; WT *Pst*: Col-0 *Pst* vs. Col-0 mock; Mut. *Pst*: *nac53-1 78-1 Pst* vs. *nac53-1 78-1* mock; Genot. *Pst*: *nac53-1 78-1 Pst* vs. Col-0 *Pst*. (H) Level of differential expression (Log2FC) of the 945 DEGs found in Genot. *Pst* (see panel G) for the several conditions analyzed. (I) Cladogram of top 40 GO terms for BP enriched in the list of 945 DEGs. Colors represent broad GO clusters. (J) Venn diagram representing the overlap between the 945 DEGs and DEGs from *rpt2a-2 Pma* (|Log2FC| > 0.5) or Col-0 MG132 24h (|Log2FC| > 0.5). (K) Cladogram of top 30 GO terms for BP enriched in the list of 538 DEGs extracted from the combined analysis (panel J). For conciseness terms are hidden and grouped in 4 processes. (L) Protein network of the genes from the 945 DEGs list related to the GO BP terms from panel K. Node colors refers to the 4 GO clusters from panel K.

### NAC53 and NAC78 coordinate the regulation of proteasome and photosynthesis gene expression through the same cis-element

The global transcriptional profile during proteotoxic stress indicates a strong link between proteasome activation and repression of photosynthesis. Thus, we hypothesized NAC53/78 can act as novel repressors of PhANGs. Detailed analysis of distinct photosynthetic processes revealed that while many processes were downregulated during proteotoxic stress, loss of NAC53/78 had a general effect on PhANGs transcriptional regulation (Extended data Fig. 5a). Enrichment analysis for specific promoter cis-elements in the 4 gene clusters of the core transcriptional network (Fig. 4k,l) revealed that the proteasome genes and PhANGs share a common cis-element characterized by the PRCE-like [TGGGC] core motif (Fig. 5a). In line with this, PRCE-like elements appeared homogeneously distributed among PhANGs promoters and enriched close to the transcription starting site (Extended data Fig. 5b,c). We then extracted the promoter regions of selected candidates, LHCa3 and PSAD1, (Fig. 5b), members of the light harvesting complex I and photosystem I^26^. Performing electromobility shift assay (EMSA), we could confirm that NAC53 and NAC78 directly bind to these elements (Fig. 5c), dependent on the presence of their respective DNA recognition motif (Fig. 5c). Furthermore, we confirmed the TGGGC motif as being the driver of this association, as mutated oligos were unable to outcompete the association of NAC53/78 with the probes (Fig. 5d). Chromatin IP followed by qPCR corroborated our findings, as both TFs were only able to associate to the selected promoter regions when plants were challenged with bacteria-induced proteotoxic stress (Fig. 5e). Performing luciferase reporter assays in protoplasts ^27^, we revealed that the PSAD1 promoter activity was consistently repressed by NAC53 and 78, which did not occur in *nac53-1 78-1* mutant protoplasts (Extended data Fig. 5d). Deletion of the PRCE motif in PSAD1 abolished NAC78-mediated repression, while NAC53 led to an increase in promoter activity (Extended data Fig. 5e). In contrast, LHCa3 promoter activity appeared to be rather induced by NAC53 and unchanged by NAC78 (Extended data Fig. 5d), with NAC53-mediated activation being dependent on the PRCE motif (Extended data Fig. 5e). This difference might be due to the fact that both TFs are expressed without a TM domain, bypassing ERAPS, in a rather artificial system. To circumvent this, we analysed the gene expression of *LHCa3* and *PSAD1* in the transgenic NAC53 and 78 lines upon BTZ treatment. Proteasome inhibition in the transgenic NAC53 and 78 lines partially enhanced the repression of PhANGs, supporting the notion that NAC53/78 act as repressors of photosynthesis during proteotoxic stress (Extended data Fig. 5f). In addition, both lines showed an increase in *PBA1* and *RPT2a* mRNA (Extended data Fig. 5g), confirming that NAC53/78 can act concomitantly as transcriptional repressor and activator. These results confirmed the importance of the PRCE-like element in PhANGs promoters, suggesting that NAC53 and NAC78 possess the ability to modulate each other’s activity and highlights the complexity of the NAC53/78-PRCE regulatory module mediating transcriptional repression of PhANGs and activation proteasome genes. To confirm that NAC53/78 act indeed as novel repressors of photosynthesis we monitored the abundance of photosynthesis proteins, PSII activity and photosynthetic pigment content upon bacteria-induced proteotoxicity. To summarize, we detected that bacterial infection had a significantly lesser impact on all measured photosynthesis readouts in the *nac53-1 78-1* double mutant compared to Col-0 (Fig. 5f-j). Consistently with these findings, subjecting the NAC53/78 transgenics to constant proteasome inhibition resulted in strong developmental defect (Extended data Fig.5h,i), suggesting that the role of NAC53/78 on photosynthesis repression is not restricted to bacterial infection. Altogether, we demonstrate the unique ability of NAC53 and NAC78 to mediate activation and repression of target genes through the same PRCE element. This makes them novel repressors of photosynthesis during proteotoxic stress and explains our previous findings that the NAC mutant was more tolerant to organelle-directed proteotoxic stress.

**Fig. 5.**
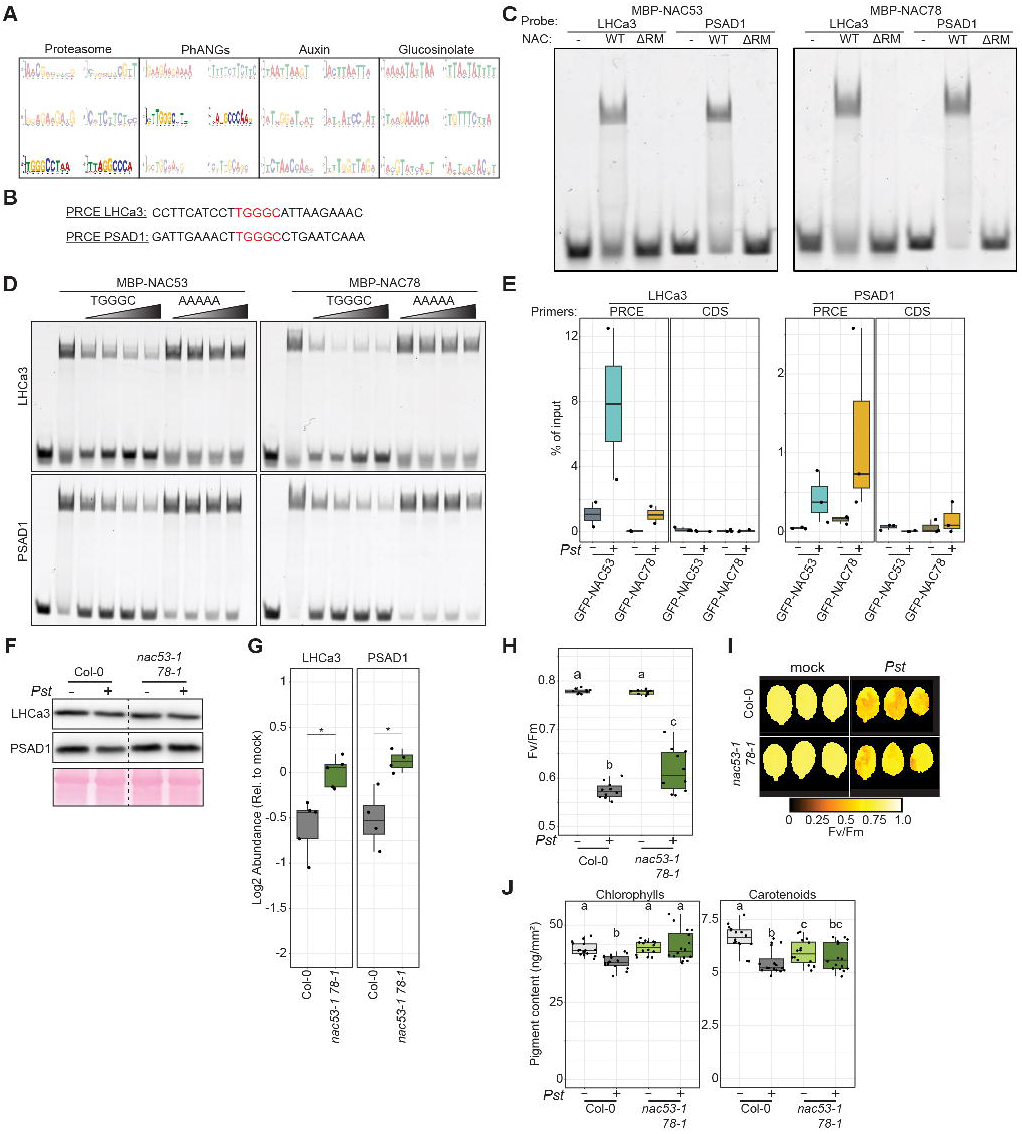
NAC53 and NAC78 coordinate the regulation of proteasome and photosynthesis gene expression through the same cis-element. (A) Top 3 motifs found by STREME software in the promoter regions from the 4 network clusters (Figure 4L). PRCE-like elements from proteasome and PhANGs clusters are highlighted. (B) 25bp region comprising the PRCE-like elements from LHCa3 and PSAD1 promoters used in this figure. The region corresponds to the probe sequence used in panel C and D. The PRCE is highlighted in red. (C) Electro-Mobility Shift Assay (EMSA) of MBP-NAC53/78 with LHCa3/PSAD1 probes (see panel B). As a negative control, probes were incubated without MBP-NAC53/78 (-) or with MBP-NAC53/78 deleted for their DNA recognition motif (ΔRM). Experiment was repeated twice with similar results. (D) EMSA competition assay of MBP-NAC53/78 association with LHCa3/PSAD1 probes. Competitor is applied at a concentration gradient (25X, 50X, 75X and 100X) indicated by the shade of grey. WT and mutated competitors are labelled TGGGC and AAAAA, respectively. Experiment was repeated twice with similar results. (E) Boxplot representing association of GFP-NAC53/78 with LHCa3/PSAD1 PRCE in adult *A. thaliana* transgenics lines infected with *Pst* or mock for 8h after Chromatin Immunoprecipitation. The association is quantified as % of input and amplification of a region in the CDS is used as negative control. (F) Immunoblot against mature PhANGs proteins in Col-0 or *nac53-1 78-1* adult plant leaves inoculated with *Pst* or mock solutions for 24h. Ponceau staining is used as loading control. (G) Relative Log2FC abundance of PhANGs mature proteins immunoblotted in panel F. The abundance is calculated relative to the mock conditions. Statistical differences are assessed via a Welch t.test (p values: n.s > 0.05, * < 0.05, ** < 0.01). (H) Photosystem II (PSII) activity measurement on Col-0 or *nac53-1 78-1* adult plants inoculated with *Pst* or mock solutions for 24h. Letters indicate the statistical group assessed by pairwise Welch t.test (p value < 0.05). The experiment was repeated 3 times with consistent results. (I) Representative pictures of measurements from panel I. Fv/Fm values are represented by false color as indicated by the color gradient legend. (J) Total chlorophylls and carotenoids from Col-0 or *nac53-1 78-1* adult plant leaves inoculated with *Pst* or mock solutions for 24h. Letters indicate the statistical group assessed by pairwise Wilcoxon-Mann-Whitney test (p value < 0.05). The experiment was repeated 3 times with consistent results.

### The proteasome autoregulatory feedback loop monitors photosynthesis homeostasis during stress responses

Our results suggest that photosynthesis repression by NAC53/78 is a general feature of the proteasome autoregulatory feedback loop to cope with proteotoxicity. In line with this, recent evidence indicates that loss of proteasome function in a chloroplast import deficient mutant improved photosynthesis due to a lack of chloroplast precursor protein degradation^28^. To analyze to which extent NAC53/78 are involved in the response to chloroplast perturbation, we investigated photosynthesis readouts upon BTZ and Lincomycin treatments in *nac53-1 78-1* plants. All readouts (pigment content, PhANGs expression, PSII activity) were significantly less impaired in *nac53-1 78-1* (Fig. 6a, b and Extended data Fig. 6a), suggesting that NAC53/78 are indeed monitoring PhANGs expression during chloroplast stress. We then repeated the phenotypic screen introduced in Fig. 1 using the *hrd1a 1b* double mutant^29^ to investigate the direct involvement of the proteasome autoregulatory feedback loop in coordinating cellular proteostasis and photosynthesis. Proteasome inhibition rendered the *hrd1a 1b* more tolerant in comparison to Col-0 (Fig. 6c), which is consistent with its role in negatively regulating NAC53/78 stability (Fig. 3g). In contrast, all other tested drugs or bacterial infection rendered the *hrd1a 1b* mutant more susceptible (Fig. 6c,d). The reduced growth of *hrd1a 1b* upon chloroplastic perturbations (Cml and Lin) are consistent with our previous results (Fig. 1b), as stabilisation of NACs would lead to increased repression of PhANGs and growth reduction (Extended data Fig. 5a, h,I). The impaired performance of *hrd1a 1b* upon CDC48 inhibition, pathogen infection or ER stress induction might be due to the general function of HRDs in ERAD^30^. Analysis of the photosystem II activity revealed a stronger reduction in the *hrd1a 1b* double mutant (Extended data Fig. 6b) upon infection corroborating our findings that ERAPS coordinates proteasome and photosynthesis through NAC53/78 stability. In addition, these data suggest that organellar perturbation would lead the NAC53/78 activation. Indeed, subjecting GFP-NACs transgenics to several organellar perturbators led to stabilization and nuclear localization of both TFs (Extended data Fig. 6c). Taken together our analysis illustrates the importance of the proteasome autoregulatory feedback loop in maintaining the fine equilibrium of proteostasis to ensure normal response to proteotoxic stress and suggests that it is involved in other biological contexts. To answer this, we performed a large-scale meta-transcriptomic analysis using the recently developed Arabidopsis RNAseq data base^31^. Using our pipeline (see methods and Supplementary Table 3) we investigated the correlation of transcript abundance between the 54 genes encoding 26S proteasome subunits and the 68 PhANGs associated with proteotoxic stress. We could observe that genes within the same cluster displayed a strong positive correlation (ρ ≈ 0.6 and ρ ≈ 0.5 for 26S Proteasome and PhANGs, respectively). Strikingly, analysing the relationship between the two clusters Proteasome vs. PhANGs showed a significant negative correlation (ρ ≈ -0.4) (Fig. 6e), supporting the notion that this is a general phenomenon. To investigate this in more detail, we extracted 43 projects in which 26S Proteasome and PhANGs appear transcriptionally co-regulated (Supplementary Table 3). From these 43 projects, 35 displayed an opposite regulation for 26S Proteasome and PhANGs mRNA level (Extended data Fig. 6d), confirming that both clusters undergo recurrent contrasting co-regulation. Amongst them pathogen attack, abiotic stress (cold, dark, drought), circadian rhythm, nutrient stress, hormone treatment (SA, auxin) as well as treatment with phytotoxins (trans-chalcone) resulted in a negative correlation. It suggests that coordination of PhANGs/proteasome is a general feature of plant stress responses and developmental processes. Thus, the coordination of proteasome activation and PhANGs repression is a key feature of the *A. thaliana* transcriptome in response to multiple biotic, abiotic, and developmental cues.

**Fig. 6.**
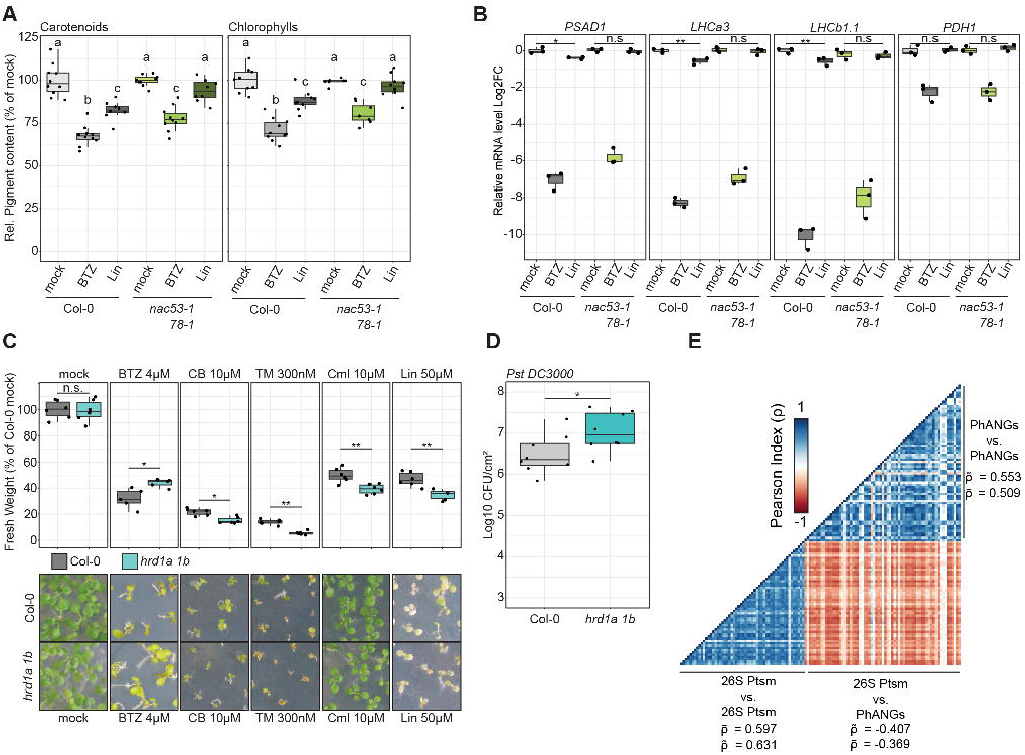
The proteasome autoregulatory feedback loop monitors photosynthesis homeostasis during stress responses. (A) Photosynthetic pigment content from Col-0 or *nac53-1 78-1* adult plant leaves infiltrated with Lin 200µM, BTZ 2µM or mock solutions for 24h. Content is expressed in percentage relative to Col-0 mock. Letters indicate the statistical group assessed by pairwise Wilcoxon-Mann-Whitney test (p value < 0.05). The experiment was repeated 3 times with consistent results. (B) Log2FC mRNA level of the indicated transcripts in Col-0, *nac53-1 78-1* adult plants leaves Lin 200µM, BTZ 2µM or mock solutions for 24h. Log2FC is calculated from Col-0 mock and *PDH1* is used as negative control. Statistical differences are assessed with a Welch t-test (p values: n.s. > 0.05, * < 0.05, ** < 0.01). The experiment was repeated twice with consistent results. (C) Fresh weight of seedlings grown under the indicated treatments at 10-14 day after germination. The fresh weight is represented as a percentage of Col-0 mock conditions. Letters indicate the statistical groups assessed by pairwise Wilcoxon-Mann-Whitney test (p value < 0.05). Every treatment has been repeated at least twice with consistent results. Representative phenotypic pictures are included. (D) Bacterial density shown as Log10 Colony-Forming-Unit (CFU) per leaf cm² in Col-0 and *hrd1a hrd1b* plants. Statistical significance is assessed by a Welch t-test (p values: * < 0.05). The experiment has been repeated three times with consistent results. (E) Correlation heatmap of the 54 26S proteasome subunit genes and the expression of 68 PhANGs from publicly available 1223 RNAseq libraries identified in this study.

## Discussion

Understanding how the proteasome can degrade proteins from different compartments and how these different subcellular signals are integrated upon external cues and to maintain overall proteostasis was a long-standing question. Our study discovered that the NAC53/78 module acts as a gatekeeper to facilitate the communication between the chloroplast-proteasome-nucleus during proteotoxic stress (Extended Data Fig. 7). We have identified ERAPS as the key regulatory node of the proteasome autoregulatory feedback loop that shapes and maintains subcellular proteostasis in plants. Considering this loop is mechanistically conserved across kingdoms, similar control mechanisms of subcellular proteostasis are possible in yeast and in animals^6^. Interestingly, connections between ER-anchored TFs and proteasome activation have previously been observed in animals, whilst separate studies identified an indirect link between proteasomal, and mitochondrial transcriptional regulation in yeast^19^. However, a unified mechanism that explains the direct crosstalk between these seemingly disparate processes has been lacking. This implies a divergence between yeast and plants on subcellular proteostasis coordination.

Our findings highlight that subcellular proteostasis is governed by the ERAPS-dependent autoregulatory pathway permitting a rapid integration of proteotoxic signals from various compartments. As such, it allows the communication between different organelles such as chloroplast or mitochondria with the nucleus, a process referred as retrograde signaling^32^. This communication is essential to respond to sudden changes in the environment. Retrograde signaling from mitochondria or chloroplast to the nucleus involves massive transcriptional changes that in turn influence organelle function^33^. Some nuclear communicators have been identified to mediate retrograde signaling^33^. However, to date, transcription factors GLK1 and 2, are the only known PhANGs activators^34^, while putative repressors ABI4 and PTM could not be verified as retrograde signaling components^35–38^. Here we have identified that plants have evolved a dual-regulatory system through the proteasome autoregulatory feedback loop: the NAC53/78-PRCE module acts as a novel retrograde signaling component, repressing PhANGs expression and activating proteasome genes via the same cis-element during proteotoxic stress. In this scenario, (i) repression of PhANGs by NAC53/78 may be a strategy to cope with the excess of precursor accumulation due to proteasomal stress (Extended Data Fig. 7) which is corroborated by our findings that the *nac53-1 78-1* mutant is more tolerant to photosynthesis stress and HSP90 inhibition which activate NAC53/78 nuclear localization but also (ii) to counteract the overactivation of PhANGs by GLK1 that is degraded by the proteasome^39^. To our knowledge, a similar system has not been found in other multicellular organisms such as animals. However, findings in animals that connect proteasome activators and mitochondrial biogenesis^40^, strongly suggest a tight link between proteoasome and energy metabolism in animals.

Various environmental perturbations impede the balance between protein translation, sorting and degradation resulting in proteotoxicity. Mounting evidence highlights a role for the proteasome in clearing organelle-associated proteins and immune components, mediating the growth-defense trade-off. Thus, it is essential to decipher how signals from different compartments and organelles are integrated to maintain proteostasis during various stress conditions. Our findings provide a new conceptual framework for understanding the integral role of the proteasome complex as a *bona fide* signalling hub, to manage cellular proteostasis under environmental stress. This framework could be potentially universally employed across various organisms to adapt to internal and external disruptions.

## FIGURE LEGENDS

**Extended data Fig. 1. Deciphering the details of NAC53/78 interactome.**

(A) Chimeric NAC53/78 constructs generated in this study. Upstream promoter, fluorescent tag and protein domain of interests are represented.

(B) Immunoblot against GFP on crude extracts of *N. benthamiana* leaves transiently expressing sTag-NACs in addition to mock or 6h BTZ (10µM) treatment. Ponceau staining is used as loading control.

(C) Immunoblot against GFP or RFP on crude extracts of *N. benthamiana* leaves transiently expressing dTag-NACs in addition to mock or BTZ 10µM for 6h. Ponceau staining is used as loading control.

(D) Immunoblot against GFP after immunoprecipitation (IP) with anti-GFP agarose beads on *N. benthamiana* leaf extracts transiently expressing sTag-NACs after mock or *PstΔHopQ* infection for 8h. The same samples were used for subsequent MS/MS analysis.

(E) Immunoblot against GFP or RFP after IP with anti-GFP or anti-RFP agarose beads on *N. benthamiana* leaf extracts transiently expressing dTag-NACs. The same samples were used for subsequent MS/MS analysis.

(F) Cladogram of top 30 GO terms for cellular components enriched in the list of *A. thaliana* ortholog proteins found in NAC53/78 common interactome.

(G) Subcellular compartments enrichment in the list of *A. thaliana* proteins from NAC53/78 common interactome based on their SUBA5db annotation. Statistical significance is provided through a chi-square testing.

(H-K) Heatmap representing the peptide enrichment (Log2FC) of the protein clusters from the network Fig. 2G.

**Extented data Fig. 2. NAC53/78 are subjected to high level of poly-ubiquitination.**

(A) *In vitro* analysis of LS-102 effect on HRD1a/b ubiquitination activity. Removal of E1 and ATP from the reaction was used as negative control.

(B) Quantification of the inhibitory effect of LS-102 (100µM) on HRD1 activity. Bar plots represent the mean of three independent replicates with error bars representing the standard deviation.

(C) Heatmaps representing the mean Log10 intensity of peptides bearing a di-glycine mark on the indicated lysin residue identified by MS/MS analysis of *in vitro* trans-ubiquitination reactions (see Figure 3F).

(D) Heatmaps representing the mean Log10 intensity of peptides bearing a di-glycine mark on the indicated lysin residue identified by MS/MS analysis of IP samples generated for Figure 2.

(E) Alignments of NAC53 and NAC78 peptide sequence regions around the identified lysin residues in panel C and D. Colors represent the residue conservation (green: conserved, shade of red: not conserved). All the identified lysins in one protein are conserved in the other. Underlined residues correspond to the mutated residues generated for the panel H and I.

(F) *In vitro* trans-ubiquitination assays of GST-HRD1a/b against MBP-NAC53/78 in the presence of LS-102 inhibitor. Removal of E1 and ATP from the reaction was used as negative control.

(G) Immunoblot analysis of *N. benthamiana* transiently expressing free-GFP, RFP-HDEL or dTag-NAC53/78 subjected to IP with anti-ubiquitin beads. The experiment was repeated twice with similar results.

(H) Immunoblot analysis of *N. benthamiana* transiently expressing GFP-NAC53/78 mutated for 5 lysins (Mut., see panel E) or not (WT) subjected to IP with anti-ubiquitin beads.

(I) Quantification of the relative ubiquitination levels assessed by the ratio between GFP signal and ubiquitin signal after IP as in panel H. Barplots represent the mean ratio of two replicates, normalized to the ratio of WT constructs with error bars representing the standard deviation.

**Extended data Fig. 3. NAC53/78 association with CDC48 is required for their nuclear translocation.**

(A) Confocal microscopy pictures of transgenic GFP-NAC53/78 root seedlings exposed to BTZ 10µM, BTZ 20µM, CB-5083 10µM or CB-5083 20µM for 3h. Scale bars = 10µm

(B) Confocal microscopy pictures of *N. benthamiana* leaves transiently co-expressing AtCDC48c-GFP with RFP-NAC53/78. Co-localization is visible in aggregate-like structures.

Scale bars = 1µm

(C) Confocal microscopy representative pictures (used for panel D-E) of transgenic GFP-NAC53/78 roots treated with BTZ 10µM and CB-5083 10µM followed by Propidium Iodide staining. Scale bars = 10µm

(D-F) Boxplot representing the GFP signal quantification after confocal microscopy imaging of transgenic GFP-NAC53/78 roots exposed to BTZ 10µM or CB-5083 10µM. Black dots represent one cell. Statistical difference is assessed by a Wilcoxon-Mann-Whitney test.

**Extended data Fig. 4. Transcriptional analysis of proteotoxic stress reveals a trade-off between 26S proteasome and other biological process.**

(A) mRNA level Log2FC of all 26S proteasome genes and associated interactors/chaperones in several conditions of the *nac53-1 78-1 Pst* RNAseq.

(B) Number of differentially expressed genes (DEGs) in the different conditions of the *nac53-1 78-1 Pst* RNAseq analysis. Total number of DEGs per condition is indicated. Colors indicate the proportion of DEGs up-regulated and down-regulated. DEGs are considered when |Log2FC| > 1.5. Genot. mock: *rpt2a-2* mock vs. Col-0 mock; WT *Pma*: Col-0 *Pma* vs. Col-0 mock; Mut. *Pma*: *rpt2a-2 Pma* vs. *nac53-1 78-1* mock; Genot. *Pma*: *rpt2a-2 Pma* vs. Col-0 *Pma*.

(C) mRNA levels Log2FC of the 981 DEGs found in Genot. *Pma* (see panel D) for the several conditions analyzed.

(D) mRNA level Log2FC of all 26S proteasome genes and associated interactors/chaperones in the several conditions of the *rpt2a-2 Pma* RNAseq.

(E) Cladogram of top 40 GO terms for BP enriched in the list of 945 DEGs. Colors represent the broad GO clusters.

(F) mRNA level Log2FC of the genes present in the 4 network modules from Figure 4L in the 3 Transcriptome used.

**Extended data Fig. 5. Characterization of the NAC53/78-PRCE module in PhANGs promoters.**

(A) mRNA level Log2FC of the PhANGs clusters from Figure 4L in the 3 transcriptomes used. Genes are separated based on their associated process in the photosynthetic pathway.

(B) Mapping of the PRCE-like motifs in PhANGs promoter regions from panel A. Promoter regions are organized according to the order of the genes from panel A. Purple marks represent the position of the motifs based on their distance from the transcription starting site (TSS). The two red arrows highlight LHCa3 and PSAD1 promoter regions.

(C) Histograms representing the number of occurrences of the PRCE-like motifs across the 4 genes cluster from the network Figure 4L. Each bar represents a 20bp region, frequency *f* of promoters bearing at least one motif is indicated as a percentage.

(D) LHCa3/PSAD1 promoter activity measured by luminescence in Col-0 or *nac53-1 78-1* protoplasts extract transiently co-expressing LHCa3/PSAD1 reporter constructs and NAC53/78ΔTM. Log2FC is calculated from the luminescence level of protoplasts expressing the reporter construct alone. Replicates are a pool of at least two independent experiments. Letters indicate the statistical group assessed by pairwise Wilcoxon-Mann-Whitney test (p value < 0.05).

(E) Promoter activity as calculated in panel D for LHCa3/PSAD1 promoters deleted for their respective DNA regions from Figure 5B. Replicates are a pool of at least two independent experiments. Letters indicate the statistical group assessed by pairwise Wilcoxon-Mann-Whitney test (p value < 0.05).

(F-G) Boxplot representing the Log2FC mRNA level the indicated transcripts in Col-0, GFP-NAC53 or GFP-NAC78 seedlings treated with BTZ 10µM or mock solutions for 6h. Log2FC is calculated from Col-0 mock. Letters indicate the statistical group assessed by pairwise Wilcoxon-Mann-Whitney test (p value < 0.05).

(H) Fresh weight of seedlings from indicated genotypes grown under the indicated treatments at 10 days after germination. Letters indicate the statistical group assessed by pairwise Wilcoxon-Mann-Whitney test (p value < 0.05). Experiment was repeated 3 times with consistent results.

(I) Representative phenotypic pictures related to panel D.

**Extended data Fig. 6. The proteasome autoregulatory feedback loop transcriptional signature is a systemic response to environmental cues.**

(A) PSII activity measurement on Col-0 and *nac53-1 78-1* treated as indicated in Figure 6A. Replicates are a pool of 3 independent experiments. Letters indicate the statistical group assessed by pairwise Wilcoxon-Mann-Whitney test (p value < 0.05).

(B) PSII activity measurement on Col-0 and *hrd1a hrd1b* adult plant leaves inoculated with *Pst* or mock solutions for 24h. Letters indicate the statistical group assessed by pairwise Welch t.test (p value < 0.05). The experiment was repeated 2 times with consistent results.

(C) Confocal microscopy pictures of transgenic GFP-NAC53/78 *A. thaliana* roots exposed to mock treatment, MV 100µM, GDA 100µM and Lin 100µM for 2h. The treatments were repeated at least three times with similar observations. Scale bars = 10µm

(D) Density functions of the 54 26S proteasome genes and 68 PhANGs transcripts (Log2FC) in the different transcriptome projects. Log2FC limits are -10 to +10. Treatment types and relevant condition information are provided in the figure. Plots are separated by treatment groups and subdivided by experiments.

**Extended data Fig. 7. ER-anchored protein sorting (ERAPS) controls the fate of two proteasome activators for intracellular organelle communication during proteotoxic stress.**

In steady state conditions, the ER-anchored protein sorting system (ERAPS) promotes the constitutive degradation of NAC53/78 via the 26S proteasome. Meanwhile, PhANGs expression is activated and subsequent chloroplastic import permits the maintenance of active photosynthesis. Upon proteotoxic stress, the chloroplast import machinery is targeted for proteasomal degradation. This leads to an accumulation of PhANGs precursors which are therefore subjected to proteasomal degradation, inducing proteotoxicity. Thus, to avoid proteotoxicity, the ERAPS system facilitates the nuclear translocation of NAC53/78, to activate the production of a new proteasome complex and to repress PhANGs expression to mitigate substrate accumulation.

## Methods

### Plant materials and growth conditions

*A. thaliana* Columbia-0 (Col-0) ecotype was considered as WT. T-DNA insertion mutants for *NAC53* (nac53-1, SALK_009578C) and *NAC78* (nac78-1, SALK_025098) were previously described in Gladman et al., 2016. T-DNA insertion mutants for *RPT2a* was previously described^20^. T-DNA insertion mutants for *HRD1a (hrd1a*, SALK_032914) and *HRD1b* (*hrd1b,* SALK_061776) were obtained from Yasin Dagdas. *promUBQ10*::GFP-NAC53 and *promUBQ10*::GFP-NAC78 transgenics lines were obtained by *Agrobacetrium tumefaciens* floral dipping of Col-0 plants. For experiments on seedlings, seeds were surface sterilized 10min with 1.3% sodium hypochlorite. Seeds were sown on ½ Murashige and Skoog (MS) medium plus 1% sucrose and stratified for 2 days. Plants were grown under long day conditions (light/dark cycles: 16h 22°C/8h 20°C, 130µmol.mm².s^-1^ light intensity, 70% relative humidity). For experiments on adult plants, plants were grown under short day conditions (light/dark cycles: 12h 22°C/ 12h 20°C, 90µmol.mm².s^-1^ light intensity, 70% relative humidity). Plants were grown 4-5 weeks until use.

### Nicotiana benthamiana growth

*N. benthamiana* plants were grown under long day conditions (light/dark cycles: 16h/8h, at 21°C and 70% humidity) and typically used 4 weeks after germination for confocal microscopy experiments and 5 weeks after germination for co-immunoprecipitation experiments.

### Bacterial strains

*For Pseudomonas syringae* pv. *tomato* DC3000 wild-type and *Pseudomonas syringae* pv. *maculicola* strains ES4326 were grown on King’s B medium containing rifampicin 100µg.mL^-1^ at 28°C. For floral dipping and transient expression in *Nicotiana benthamiana* leaves, *Agrobacterium tumefaciens* C58C1 strain was grown in LB broth high salt (Duchefa L1704) containing the required antibiotics at 28°C. For molecular cloning, *Escherichia coli* Top10 strain was grown in LB broth high salt (Duchefa L1704) containing the required antibiotics at 37°C. For protein purification, *Escherichia coli* BL21 (DE3) was grown in LB broth high salt (Duchefa L1704) containing the required antibiotics at 37°C or 16°C.

### Gene Accession

Arabidopsis genome initiative (AGI) locus identifier of the principal genes investigated in this study are the following: *NAC53* (AT3G10500), *NAC78* (AT5G04410), *HRD1a* (AT3G16090), *HRD1b* (AT1G65040), *CDC48c* (AT5G03340), *RPT2a* (AT4G29040), *LHCa3* (AT1G61520), *PSAD1* (AT4G02770).

### Molecular cloning

For chimeric protein constructs generated in this study, protein coding sequence (CDS) of the desired genes were amplified from *A. thaliana* cDNA. For Golden Gate based cloning^41^, CDS were amplified with addition of flanking regions including BpiI/BsaI sites and final constructs were generated using the LI CDS module of interest and desired other modules and assembled in the LIIa F 1-2 vector. For GATEWAY™ based cloning, CDS were amplified with addition of attb1/attb2 flanking regions and final constructs were generated through BP clonase™ II enzymatic reaction and LR Clonase™ II enzymatic reaction into the desired destination vectors. All LI and GATEWAY™ entry clones were verified by sanger sequencing.

### Seedlings phenotyping

For phenotyping assay, seeds were sown on ½ MS 1% sucrose round petri dish supplemented with the indicated concentration of drug or a mock treatment. After 10 to 14 days of growth fresh weight was measured. For one replicate, 5 representative seedlings were scaled together on an analytical scale (resolution 0.0001g). Typically, 5 replicates were measured for subsequent statistical analyses.

### Transient expression in *N. benthamiana* leaves

Bacterial suspension was centrifuged for 4min at 4000g after overnight incubation of *A. tumefaciens* strain carrying the desired expression construct. Supernatant was removed and pellet was resuspended in 500µl of agroinfiltration buffer (10mM MgCl_2_, 10mM MES pH 5.7). OD600 was measured and infiltration solutions were diluted to reach final 0.5 OD600 in agroinfiltration buffer supplemented with 200µM acetoseryringone. Solution were incubated in dark at least 1h prior to infiltration. Infiltrated tissues were used 30h post infiltration for subsequent experimental procedures.

### Protein purification

Recombinant proteins were expressed in E. coli BL21(DE3) by IPTG induction. Bacterial solution was centrifuged 20min at max speed and pellet was subjected to sonication after resuspension in 1mL MBP-buffer (20 mM TRIS-HCl pH 7.5, 200mM NaC, 1mM EDTA) or 1mL IPP50 (10 mM Tris-HCl, pH 8.0. 150mM NaCl, 0.1% NP40). Sonicated samples were used for purification by affinity chromatography using amylose resin (New England Biolabs) for MBP-NAC53/78 or glutathione sepharose 4B (cytiva) for GST-HRD1a/b. For recombinant His-UBA1 and His-UBC8 protein were purified using Ni-Ted resin (Macherey-Nagel).

### *In vitro* ubiquitination assay

In vitro ubiquitination assay was performed as described previously^42^. Samples were separated by SDS–PAGE electrophoresis using 4–15% Mini-PROTEAN® TGX™ Precast Protein Gels (BioRad) followed by detection of the ubiquitinated substrate by immunoblotting using anti-MBP (New England Biolabs), anti-GST and anti-ubiquitin (Santa Cruz Biotechnology) antibodies.

### Co-immunoprecipitation

Plant tissue was homogenized in a mortar with liquid nitrogen to keep the sample frozen during the process. And extracted in 1ml/g of extraction buffer (10% glycerol, 25 mM Tris pH 7.5, 1 mM EDTA, 150 mM NaCl, 1 mM DTT, 0.5% Triton X, 1X protease cocktail inhibitor (Sigma)). Solution was vortexed for 3min and incubated in cold room on rotator for 10min following a 30min centrifugation at 4000g, 4°C. Supernatant was filtered. 50µl of filtrate was sampled as input, supplemented with 12.5µL laemmli buffer 4X (Biorad) and boiled for 10min at 95°C. Next, 10µl/g of ChromoTek GFP-Trap® or RFP-Trap Agarose beads were added and tubes were incubated 2-3h in cold room. Tubes were then centrifuge at 800g for 1min; supernatant was carefully removed and pelleted beads were transferred to 1.5mL microcentrifuge tube in 1mL of washing buffer (10% glycerol, 25 mM Tris pH 7.5, 1 mM EDTA, 150 mM NaCl, 0.5% Triton X-100, 1X protease cocktail inhibitor (Sigma)). Washing was performed by centrifugation at 800g for 1min using 1mL washing buffer. After last washing, Laemmli buffer 2X (Biorad) was added to equal amounts of beads and sample was boiled 10min at 95°C.

### RNA isolation & RT-qPCR

Total RNA isolation was performed using RNeasy Plant Mini Kit (Qiagen 74904) according to manufacturer instructions. To exclude potential contaminant DNA, RNA samples were subjected to DNase I treatment (Thermo scientific™) following provider instructions. For RNA sequencing, integrity and RNA concentration was determined (2100 Bioanalyzer, Agilent). For RT-qPCR sample analysis was performed as described previously^43^; cDNA synthesis was performed using LunaScript® RT SuperMix Kit (NEB) following provider recommendation. qPCR was performed using MESA BLUE qPCR 2X MasterMix Plus for SYBR® Assay (Eurogentec) using a 2-step reaction protocol for 40cycles with systematic evaluation of primer melting curve. mRNA level was quantified based on the ΔΔCt method followed by Log2 transformation.

### NanoLC-MS/MS analysis and data processing

For purification, proteins were subjected to a NuPAGE 12% gel (Invitrogen) and in-gel trypsin digestion was done on Coomassie-stained gel pieces with a modification: *chloroacetamide* was used instead of iodoacetamide for carbamidomethylation of cysteine residues to avoid formation of lysine modifications isobaric to two glycine residues left on ubiquitinylated lysine after tryptic digestion. Next, peptides mixture were desalted using C18 Stage tips and run on an Easy-nLC 1200 system coupled to a Q Exactive HF mass spectrometer (both Thermo Fisher Scientific) as described elsewhere^44^ with some modifications: separation of the peptide mixtures was done using a 87-min or 127-min segmented gradient from 10-33-50-90% of HPLC solvent B (80% acetonitrile in 0.1% formic acid) in HPLC solvent A (0.1% formic acid) at a flow rate of 200 nl/min. The seven most intense precursor ions were sequentially fragmented in each scan cycle using higher energy collisional dissociation (HCD) fragmentation. In all measurements, sequenced precursor masses were excluded from further selection for 30 s. The target values were 10^5^ charges for MS/MS fragmentation and 3 × 10^6^ charges for the MS scan.

Acquired MS spectra were processed with MaxQuant software package version 1.5.2.8 with an integrated Andromeda search engine. For *in vivo* IP samples, database search was performed against a *N. benthamiana* database containing 74,802 protein entries^45^, the sequences of eGFP-NAC53/78, mCerrulean3-NAC53/78-mCherry and 285 commonly observed contaminants.

For *in vitro* ubiquitination samples, database search was performed against a Uniprot *E. coli* database (4,403 entries<u>, downloaded on 7th of October 2020</u>), the sequences of MBP-NAC53/78, GST-HRD1a/b and 285 commonly observed contaminants.

Endoprotease trypsin was defined as a protease with a maximum of two missed cleavages. Oxidation of methionine, phosphorylation of serine, threonine, and tyrosine, GlyGly dipeptide on lysine residues, and N-terminal acetylation were specified as variable modifications. Carbamidomethylation on cysteine was set as a fixed modification. Initial maximum allowed mass tolerance was set to 4.5 parts per million (ppm) for precursor ions and 20 ppm for fragment ions. Peptide, protein, and modification site identifications were reported at a false discovery rate (FDR) of 0.01, estimated by the target-decoy approach (Elias and Gygi). The iBAQ (Intensity Based Absolute Quantification) and LFQ (Label-Free Quantification) algorithms were enabled, as was the “match between runs” option^46^.

### Photosynthesis monitoring

For photosynthetic pigment concentration, 2 leaf discs (6mm) were sampled and incubated over-night in 1mL acetone 100% on a rotating wheel in cold room. Tubes were next centrifuged for 3min at 400g. 200µl of was then pipetted in a transparent 96-well plate and absorbance was measured at 470nm, 646nm and 663nm using a plate reader (Tecan Infinite 200 PRO®). Acetone 100% was used as blank. Concentration was calculated as following the obtained from Karlsruher Institut für Technologie (KIT, https://www.jkip.kit.edu/molbio/998.php, “Chlorophyll and Carotenoid determination in leaves”). Values were then then expressed as ng.mm².For photosynthetic efficiency, photosystem II (PSII) activity was measured by quantifying the maximum photosystem II yield, Fv/Fm via the saturation pulse methods on dark acclimated plants (Schreiber 2004), a Imaging-PAM chlorophyll fluorometers: Maxi version v2-46i, for measurements or Dual-PAM100 (Walz GmbH, Effeltrich, Germany).

### Bacterial Infection

*Pst* overnight liquid culture was centrifuged for 15min at 4000g. Bacterial pellet was resuspended in 5mL of MgCl_2_ 10mM prior to OD600 measurement.

For qPCR, RNAseq and western blot analysis; a *Pst* solution at OD600=0.2 in 10mM MgCl_2_ was used. Adult plant leaves were syringe infiltrated with the *Pst* solution or a 10mM MgCl_2_ solution for 24h until sampling. For IP and Chromatin-IP on *A. thaliana*, *Pst* infiltration was prepared as explained above and detached whole rosettes of adult plants were vacuum infiltrated 2 times 2 min at < 15mbar. For mock, control solution (10mM MgCl_2_) was used similarly. Rosettes were put back in the growth chamber for 8h on multiple layers of wet tissue. For assessment of bacterial density, *Pst* infiltration solution was prepared in 10mM MgCl_2_ at an OD600 of 0.0001. Adult plant leaves were syringe infiltrated with the *Pst* infiltration solution for 72h in high humidity conditions. Next, 2 leaf disks per replicates were sampled and homogenized in 200µl 10mM MgCl_2_ and a serial dilution from 10^-1^ to 10^-6^ in 200µL was done. For bacterial counting, 20µL of 10^-5^ and 10^-6^ dilutions were plated. Growing colony-forming units were counted and Log10CFU/cm² was calculated.

### Chromatin Immuno-Precipitation

For Chromatin-IP, after Arabidopsis rosettes were vacuum infiltrated for 30min with a fixation buffer solution (1% formaldehyde, 10mM KH2PO4 pH 7, 50mM NaCl, 0.1M sucrose, 0.01% Triton X-100). Plant material (1g) was dried and frozen in liquid nitrogen. Tissue was ground in 5mL nuclei isolation buffer (20mM Hepes pH8, 250mM sucrose, 1 mM MgCl_2_, 5mM KCl, 40% Glycerol, 0.25% Triton X-100, 0.1mM PMSF, 0.1% 2-mercaptoethanol, 1µM BTZ). The solution was filtered and subjected to 10min centrifugation at 4°C, 3000g. Pellet was resuspended in 1mL nuclei isolation buffer and centrifuged 5min, at 4°C, 3000g, two. Pellet was resuspended in 200µl M3 buffer (10mM KH2PO4 pH7, 0.1mM NaCl, 10mM 2-mercaptoethanol) and centrifuged 5min, 4°C, 3000g. Nuclei pellet was resuspended in 1mL sonication buffer (10mM KH2PO4 pH7, 0.1mM NaCl, 0.5% Sarkosyl, 10mM EDTA). Sonication was performed with an Active Motif Sonicator (Amp 25%; Pulse: 30sec ON – 30sec OFF; Timer: 300sec). Sonicated sample was centrifuged for 5min, 4°C at max speed. Supernatant was tranferred to a new 1 tube and 10% (v/v) Triton X-100 was added to neutralize the sarkosyl. 5% (v/v) of this solution was saved as input sample. For IP, 15µL ChromoTek GFP-Trap® were activates in 15µL IP buffer (50mM Hepes pH 7.5, 150mM NaCl, 5mM MgCl_2_, 10uM ZnSO4, 1% Triton X -100, 0.05% SDS), added to the tube and incubated 3h on a rotating wheel at 4°C. Beads were washed 4 times by centrifuging for 30sec, at 1000g and subsequent incubation on rotating wheel in 500µL buffer for 3 min; twice in IP buffer, once in LNDET buffer (0.25 LiCl, 1% NP-40, 1% deoxycholate, 1mM EDTA pH 8, 10mM Tris pH 8) and once in TE buffer (10mM Tris pH 8, 1mM EDTA pH 8). After the last washing step, beads were centrifuged for 30sec at 1000g, resuspended in 200µL EB buffer (50mM Tris pH 8, 10mM EDTA pH 8, 1% SDS) supplemented with 8µL 5M NaCl to reverse cross-linking and incubated at 65°C for at least 6h. At this step input sample was added and treated similarly. For protein digestion, 1µL of Proteinase K (Thermo Scientific) was added and incubated for 1h at 45°C. Finally, DNA was purified.

For DNA quantification, qPCR was performed using MESA BLUE qPCR 2X MasterMix Plus for SYBR® Assay (Eurogentec) in technical triplicates. For quantification, % of input was calculated.

### Immunoblot analysis

For immunoblot analysis, sample processing was performed as described previously^43^.

### Electromobility Shift Assay

For EMSA assays, complementary DNA oligos were synthesized by EuroFins Genomics with ATTO565 dye linked to the 5’ end of minus strand for probes. To generate double stranded probes, complementary oligos were incubated at 2µM in annealing buffer (25mM HEPES-KOH pH 7.8, 40mM KCl) at 70°C for 5min and cool down to room temperature.

EMSA reactions were performed in 20µL reaction buffer (25 mM HEPES-KOH pH 7.8, 40 mM KCl, 1 mM DTT, 10% Glycerol) using 1µg purified MBP-NAC53/78 and 50ng DNA probe for 30min at 25°C. Reactions were subjected to native polyacrylamide migration with TGX FastCast 7.5% Acrylamide gels (Biorad) in TAE (Tris-base 40 mM pH 8.3, 20mM acetic acid, 1mM EDTA). After migration, probes were imaged using a Typhoon FLA 9000 Gel Scanner.

### Luciferase reporter assay in Protoplasts

For promoter activity analysis, *Arabidopsis* mesophyll protoplasts were generated as described perviously^27^. Luciferase quantification was performed according to Dual-Luciferase® Reporter Assay System from Promega recommendation (see <u>link</u>).

### Data analysis

For all statistical analysis, R programming language was used in the Rstudio environment. For non-omics data, the statistical test and associated p value threshold for each analysis are indicated in the figure legends. Data dispersion, medians and quantiles are represented with boxplots and replicates displayed in every figures. For bar plots in Extended Data Fig. 2, number of replicates are indicated in the figure legend, error bars represent the standard deviation from the mean.

### RNA sequencing

For *rpt2a-2 Pma* transcriptome, sequencing was performed by ATLAS Biolabs, Berlin. For *nac53-1 78-1 Pst* transcriptome, sequencing was performed by the NGS Competence Center, Tübingen. After quality control, reads were mapped to the *A. thaliana* TAIR10 reference genome. The mapped reads were counted with *htseq-count* for subsequent analysis. Log2FC and false discovery rate (FDR) were determined using R package DEseq2^47^ with default settings. A gene is considered significantly differentially expressed when its FDR < 0.05. Log2FC threshold is indicated in the figure legends according to the analysis.

### IP-MS/MS analysis

For IP-MS/MS analysis, Log2FC and false discovery rate (FDR) for each protein group were determined as described previously^42^. For downstream analysis, to each protein group a unique Agi-code was assigned by blasting every *N. benthamiana* protein present in the protein group against *A. thaliana* proteome and taking the most frequent Agi-code with the best E-value. An E-value threshold of e-10 was used, in case of absence of *A. thaliana* the protein group was not considered for downstream analysis. For heatmap representation, in case multiple protein groups correspond to the same *A. thaliana* orthologs the mean Log2FC is represented.

### Meta-Transcriptomic analysis

For meta-transcriptomic analysis in Fig. 6 and Extended data Fig. 6, fragments per kilobase per million counts were retrieved from the data base (http://ipf.sustech.edu.cn/pub/athrna/). Log2FC for individual genes in every project was calculated based on the control RNAseq libraries as annotated in the original publication^31^. Filtering of relevant projects for subsequent correlation analysis was done based on the median |Log2FC|. Median|Log2FC| > 0.5 was used for identification of projects with at least one gene cluster transcriptionally impacted and median |Log2FC| > 0.3219 was used for identification of projects in which 26S proteasome and PhANGs clusters were co-transcriptionally impacted. Pearson correlations were calculated. Only correlations with p-value < 0.05 were considered significant and used for calculation of mean/median.

### Omics downstream analysis: Gene ontology, Protein Network, subcellular enrichment & cis-element analysis

For Gene Ontology enrichment, the list of Agi-code was uploaded into ShinyGO software (http://bioinformatics.sdstate.edu/go/). Cladogram of the indicated number of top terms for the indicated type of terms was used for graphical representation. Table of annotations was extracted for further analysis.

For network creation, list of Agi-code was provided to the string database (https://string-db.org/). Table of interactions was extracted and uploaded to cytoscape software (https://cytoscape.org/) for network generation, design, and annotation. For purposes of clarity, network nodes were manually arranged. Edge transparency relate to the strength of interaction with opaque edges corresponding to strong interactions and transparent edges to weak interactions. Node color corresponds to manual annotation inferred based on GO annotations and further curated from the literature.

For subcellular compartment enrichment, the list of Agi-code was uploaded into SUBA5db (https://suba.live/). Compartment annotation was extracted, and occurrence of every compartment was counted to estimate the list distribution. For theorical distribution 20 Agi-code lists of the same length were randomly generated and processed similarly. The mean distribution of the 20 lists was used as theorical distribution. Enrichment for every compartment was calculated as a Log2FC of the ratio between the observed proportion and the theorical one.

### Cis-Regulatory Element analysis

For cis-regulatory element analysis, promoter sequences of 1000bp upstream of the transcription starting site were retrieved from plant ensembl database using biomart browser (https://plants.ensembl.org/biomart/). For identification of novel cis-regulatory elements in Figure 5A, promoter sequences of the several gene clusters were analyzed using the STREME software (https://meme-suite.org/meme/tools/streme) with default settings. For mapping and counting, promoter sequences were analyzed in R.

### Immunoblot semi-quantitative analysis

Raw image of the immublot signals against the protein of interest and the corresponding loading control were processed using Fiji software (https://imagej.net/software/fiji/downloads). Image was converted to 8-bit format, background was subtracted. Individual lanes were quantified by measurement of the peak area after circumscription with the rectangular ROI selection and “Gels” function. The obtained values for individual bands were normalized by the corresponding loading control values. Log2FC was calculated using the corresponding mock sample of the same genotype.

### Confocal microscopy imaging

Confocal microscopy was done using an inverted Zeiss LSM 880 microscope, an upright Leica SP8 microscope or an inverted Leica Stellaris 8 microscope. For mCerrulean3 imaging, excitation = 458nm and emission window = 465nm-500nm; for eGFP or sGFP imaging, excitation = 488nm and emission window = 510nm-540nm; for mRFP or mCherry imaging, excitation = 561nm and emission window = 590nm-630nm; in addition, chlorophyll auto fluorescence was vizualized in the far-red wavelength. Images were acquired with a 40X water immersion objective, pinhole set to 1 airy unit, resolution of acquisition >1024×1024 with a line average of 4.

After acquisition images were processed in Fiji Software. For images obtained from Leica microsystem devices, a gaussian blur (radius 0.75) was applied. For clarity purposes, contrast on the individual channels was manually adjusted.

For subcellular quantification of GFP signal, all images used were acquired on the Zeiss LSM 880 with a 16bits depth. Prior to imaging seedlings were incubated for 10min in a 10µM propidium iodide (PI) ½ MS solution followed by a brief washing in clear ½ MS. For PI fluorescence, was excitation = 488nm and emission window = 590nm-660nm. Each cell and corresponding nucleus were segmented using the polygon ROI selection tool in Fiji software. Signal proportion in the nucleus was quantified by dividing integrated density of nucleus ROI by integrated density of whole cell ROI and ratio of signal between nucleus and cytosol was calculated dividing nucleus ROI mean intensity by cytosolic ROI mean intensity.

For calculation of co-localization index the image J plugin “co-localization finder” was used to calculate pearson index or plot profile of GFP and RFP signals was extracted at the indicated line using Fiji software and the pearson index was calculated between the two profiles.

## RESOURCE AVALAIBILITY

Source code of R and Python3 scripts used for the several computational analysis done in this study will be put available on https://github.com/Gogz31 or can be directly requested.

## DATA AVAILABILITY

The mass spectrometry data from this publication will be made available on the PRIDE archive (https://www.ebi.ac.uk/pride/archive/) and with the identifier (PXDXXXXX; and (PXDXXXX). All relevant proteomics data are made available in the supplemental information.

## Acknowledgements

We thank Yasin Dagdas and Marja Timmermans for critical discussions during the thesis advisory committee meeting. We are thankful to #theustunlab members Ophélie Léger, Paul Gouguet for stimulating discussions and Manuel González Fuente for editorial issues. We thank Nan Wang for technical advice on the Ch-IP procedure, Robert Morbitzer for technical support on the golden gate molecular cloning. We are grateful to Ana Gabriela Andrade-Galan and Lars Pospisil for technical support on the Photosystem II activity measurement. Stefan Bieker for the help with the computational analysis. We thank Silke Wahl, Anke Biedermann and Irina Droste-Borel for their technical support in sample preparation for MS. We are thankful to Jos Schippers, Yasin Dagdas and Richard Vierstra for sharing published material. This work was supported by an Emmy Noether Fellowship GZ: UE188/2-1 from the Deutsche Forschungsgemeinschaft (DFG; to S.Ü.) and through the collaborative research council 1101 (SFB1101; to G.L.). We thank the confocal microscopy facility of ZMBP that is supported by funds from DFG (INST 37/819-1 FUGG and INST 37/965-1 FUGG) and the confocal microscopy facility of RUB (DFG Project number 523980288, GZ:INST 213/1180-1 FUGG).

## Author Contribution

S.Ü. conceived the project. G.L. performed most experiments, analyzed the data, and performed all computational analysis. M.R. performed the *in vitro* ubiquitination assays. D.B performed the EMSA, assisted with the Luciferase reporter assay and the molecular cloning. D.S. generated the *rpt2a-2 Pma* transcriptome under the supervision of F.B..M.F.W and B.M. performed LC-MS/MS analysis. S.Ü. wrote the manuscript together with G.L..

