## Extended data Fig. 1-7 for "ER-anchored protein sorting controls the fate of two proteasome activators for intracellular organelle communication during proteotoxic stress"

### SUPPORTING INFORMATION

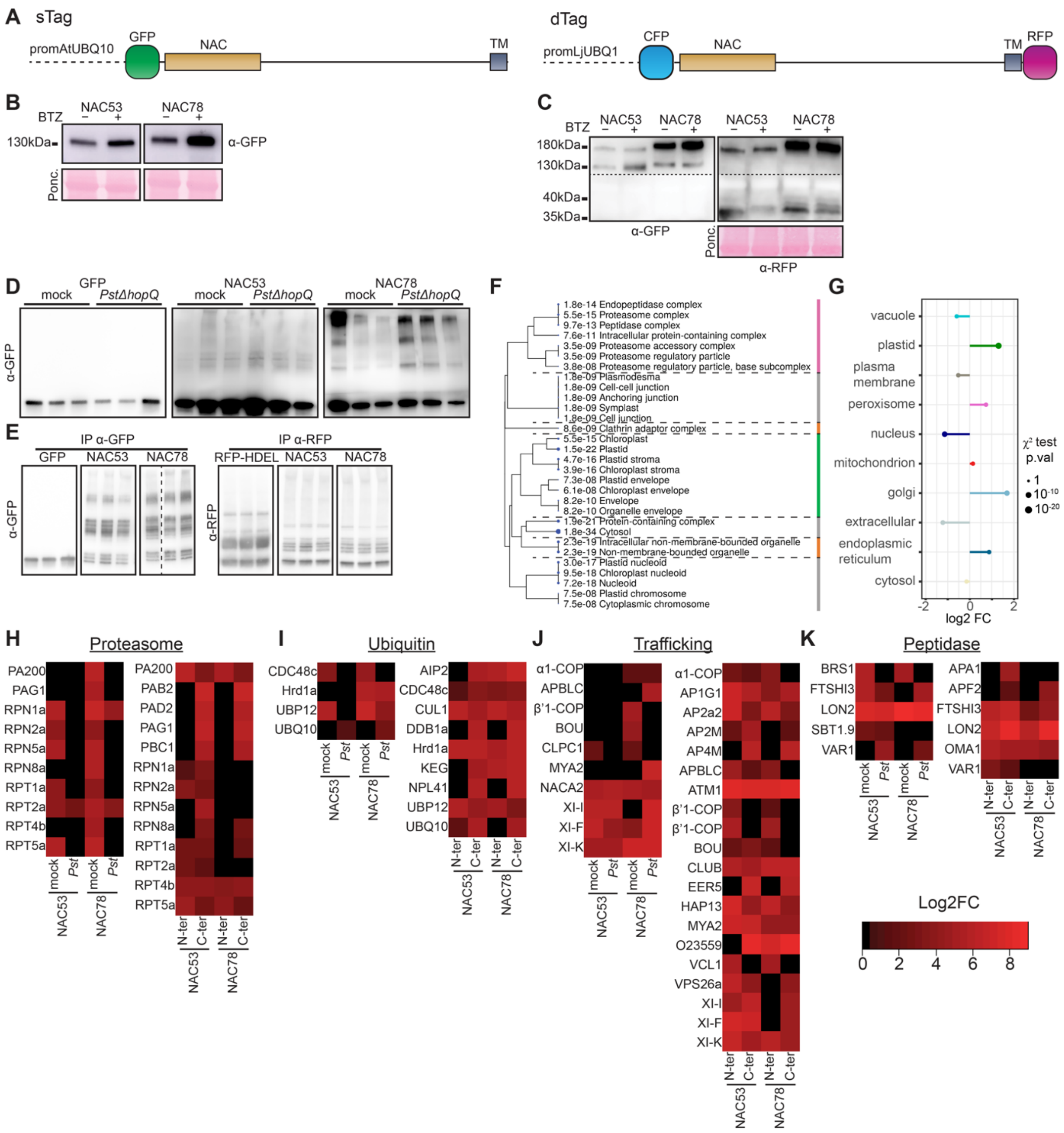


**Extended Data Fig 1. Deciphering the details of NAC53/78 interactome.**
(A) Chimeric NAC53/78 constructs generated in this study. Upstream promoter, fluorescent tag and protein domain of interests are represented.
(B) Immunoblot against GFP on crude extracts of *N. benthamiana* leaves transiently expressing sTag-NACs in addition to mock or 6h BTZ (10µM) treatment. Ponceau staining is used as loading control.
(C) Immunoblot against GFP or RFP on crude extracts of *N. benthamiana* leaves transiently expressing dTag-NACs in addition to mock or BTZ 10µM for 6h. Ponceau staining is used as loading control.
(D) Immunoblot against GFP after immunoprecipitation (IP) with anti-GFP agarose beads on *N. benthamiana* leaf extracts transiently expressing sTag-NACs after mock or *PstΔHopQ* infection for 8h. The same samples were used for subsequent MS/MS analysis.
(E) Immunoblot against GFP or RFP after IP with anti-GFP or anti-RFP agarose beads on *N. benthamiana* leaf extracts transiently expressing dTag-NACs. The same samples were used for subsequent MS/MS analysis.
(F) Cladogram of top 30 GO terms for cellular components enriched in the list of *A. thaliana* ortholog proteins found in NAC53/78 common interactome.
(G) Subcellular compartments enrichment in the list of *A. thaliana* proteins from NAC53/78 common interactome based on their SUBA5db annotation. Statistical significance is provided through a chi-square testing.

(H-K) Heatmap representing the peptide enrichment (Log2FC) of the protein clusters from the network Figure 2G.

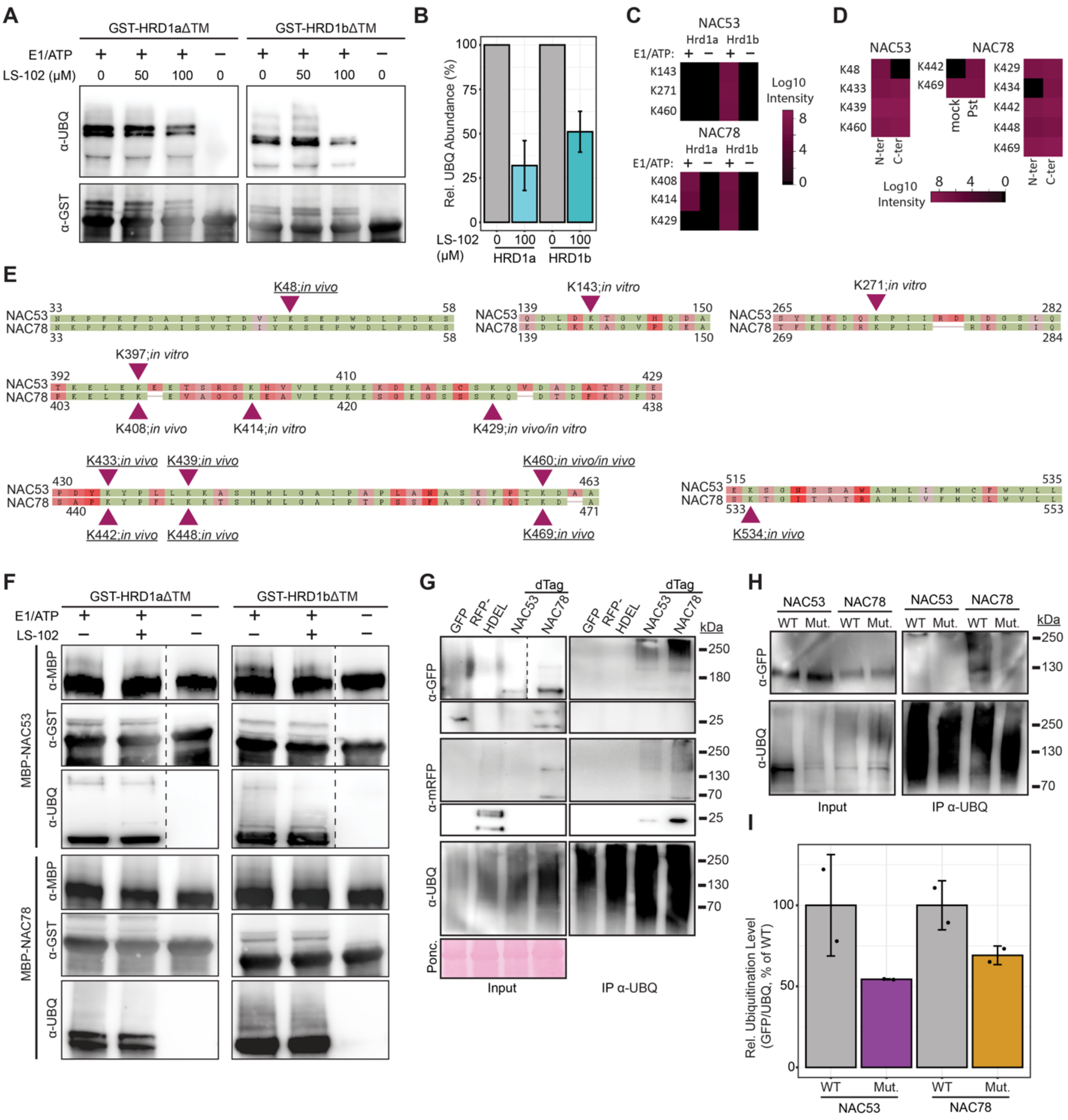

**Extended data Fig. 2. NAC53/78 are subjected to high level of poly-ubiquitination.**
(A) *In vitro* analysis of LS-102 effect on HRD1a/b ubiquitination activity. Removal of E1 and ATP from the reaction was used as negative control.
(B) Quantification of the inhibitory effect of LS-102 (100µM) on HRD1 activity. Bar plots represent the mean of three independent replicates with error bars representing the standard deviation.
(C) Heatmaps representing the mean Log10 intensity of peptides bearing a di-glycine mark on the indicated lysin residue identified by MS/MS analysis of *in vitro* trans-ubiquitination reactions (see Figure 3F).
(D) Heatmaps representing the mean Log10 intensity of peptides bearing a di-glycine mark on the indicated lysin residue identified by MS/MS analysis of IP samples generated for Figure 2.
(E) Alignments of NAC53 and NAC78 peptide sequence regions around the identified lysin residues in panel C and D. Colors represent the residue conservation (green: conserved, shade of red: not conserved). All the identified lysins in one protein are conserved in the other. Underlined residues correspond to the mutated residues generated for the panel H and I.
(F) *In vitro* trans-ubiquitination assays of GST-HRD1a/b against MBP-NAC53/78 in the presence of LS-102 inhibitor. Removal of E1 and ATP from the reaction was used as negative control.

(G) Immunoblot analysis of *N. benthamiana* transiently expressing free-GFP, RFP-HDEL or dTag-NAC53/78 subjected to IP with anti-ubiquitin beads. The experiment was repeated twice with similar results.
(H) Immunoblot analysis of *N. benthamiana* transiently expressing GFP-NAC53/78 mutated for 5 lysins (Mut., see panel E) or not (WT) subjected to IP with anti- ubiquitin beads.
(I) Quantification of the relative ubiquitination levels assessed by the ratio between GFP signal and ubiquitin signal after IP as in panel H. Barplots represent the mean ratio of two replicates, normalized to the ratio of WT constructs with error bars representing the standard deviation.

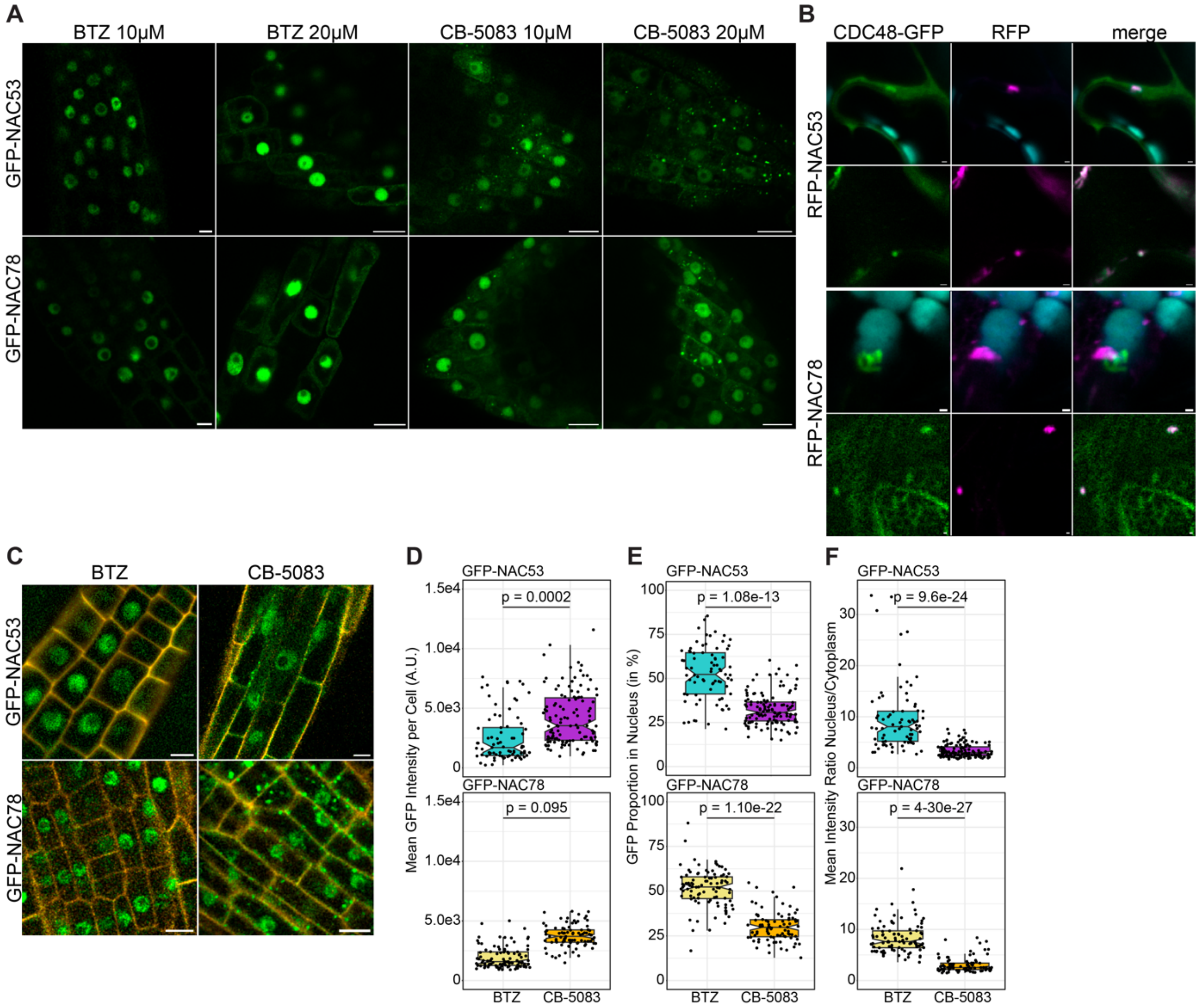


**Extended data Fig.3. NAC53/78 association with CDC48 is required for their nuclear translocation.**
(A) Confocal microscopy pictures of transgenic GFP-NAC53/78 root seedlings exposed to BTZ 10µM, BTZ 20µM, CB-5083 10µM or CB-5083 20µM for 3h.
(B) Confocal microscopy pictures of *N. benthamiana* leaves transiently co-expressing AtCDC48c-GFP with RFP-NAC53/78. Co-localization is visible in aggregate-like structures.
(C) Confocal microscopy representative pictures (used for panel D-E) of transgenic GFP-NAC53/78 roots treated with BTZ 10µM and CB-5083 10µM followed by Propidium Iodide staining.
(D-F) Boxplot representing the GFP signal quantification after confocal microscopy imaging of transgenic GFP-NAC53/78 roots exposed to BTZ 10µM or CB-5083 10µM. Black dots represent one cell. Statistical difference is assessed by a Wilcoxon-Mann-Whitney test.

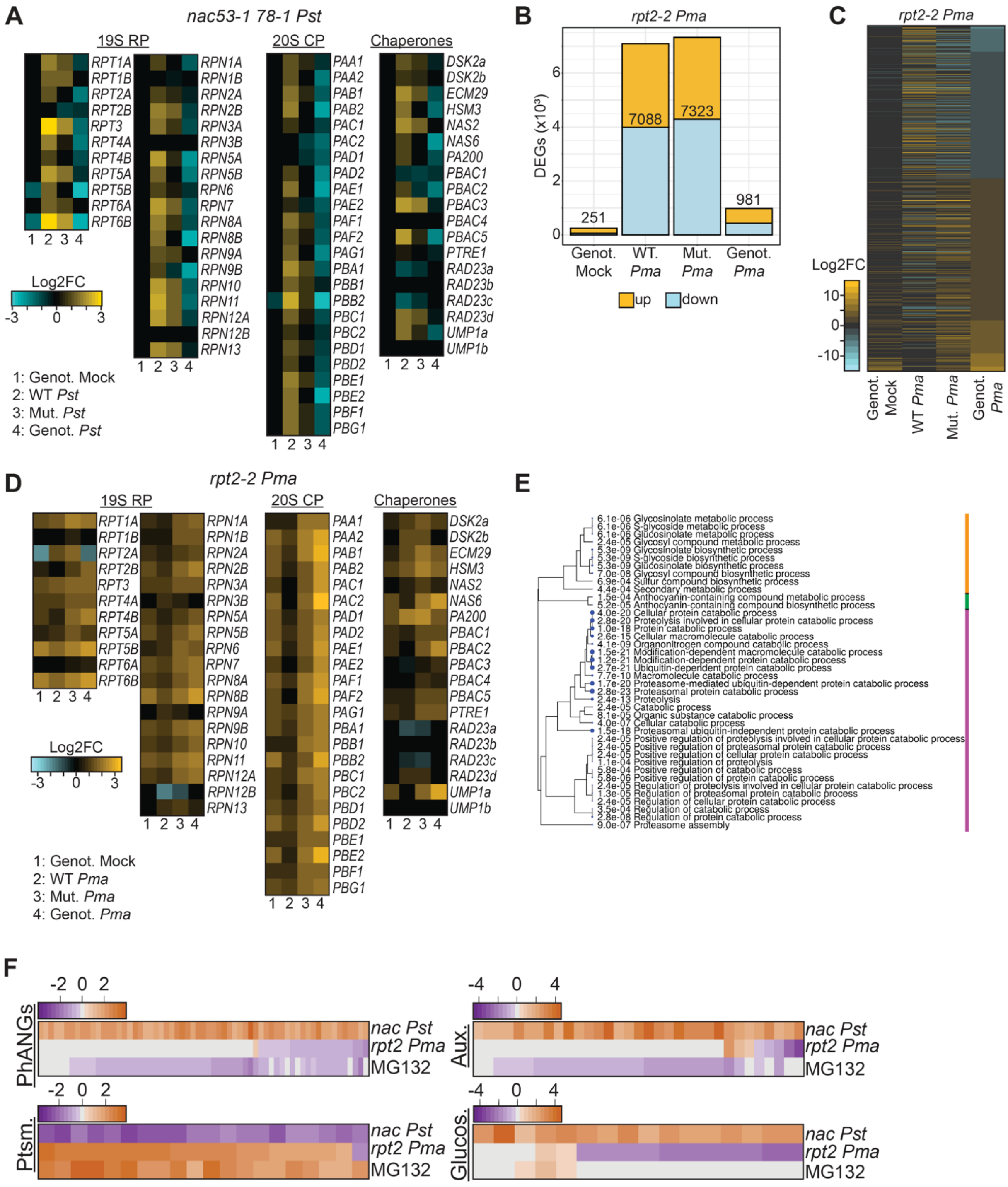


**Extended data Fig.4. Transcriptional analysis of proteotoxic stress reveals a trade-off between 26S proteasome and other biological process. Related to Figure 4**

(A) mRNA level Log2FC of all 26S proteasome genes and associated interactors/chaperones in several conditions of the *nac53-1 78-1 Pst* RNAseq.

(B) Number of differentially expressed genes (DEGs) in the different conditions of the *nac53-1 78-1 Pst* RNAseq analysis. Total number of DEGs per condition is indicated. Colors indicate the proportion of DEGs up-regulated and down-regulated. DEGs are considered when |Log2FC| > 1.5. Genot. mock: *rpt2a-2* mock vs. Col-0 mock; WT *Pma*: Col-0 *Pma* vs. Col-0 mock; Mut. *Pma*: *rpt2a-2 Pma* vs. *nac53-1 78-1* mock; Genot. *Pma*: *rpt2a-2 Pma* vs. Col-0 *Pma*.

(C) mRNA levels Log2FC of the 981 DEGs found in Genot. *Pma* (see panel D) for the several conditions analyzed.

(D) mRNA level Log2FC of all 26S proteasome genes and associated interactors/chaperones in the several conditions of the *rpt2a-2 Pma* RNAseq.

(E) Cladogram of top 40 GO terms for BP enriched in the list of 945 DEGs. Colors represent the broad GO clusters.

(F) mRNA level Log2FC of the genes present in the 4 network modules from Figure 4L in the 3 Transcriptome used.

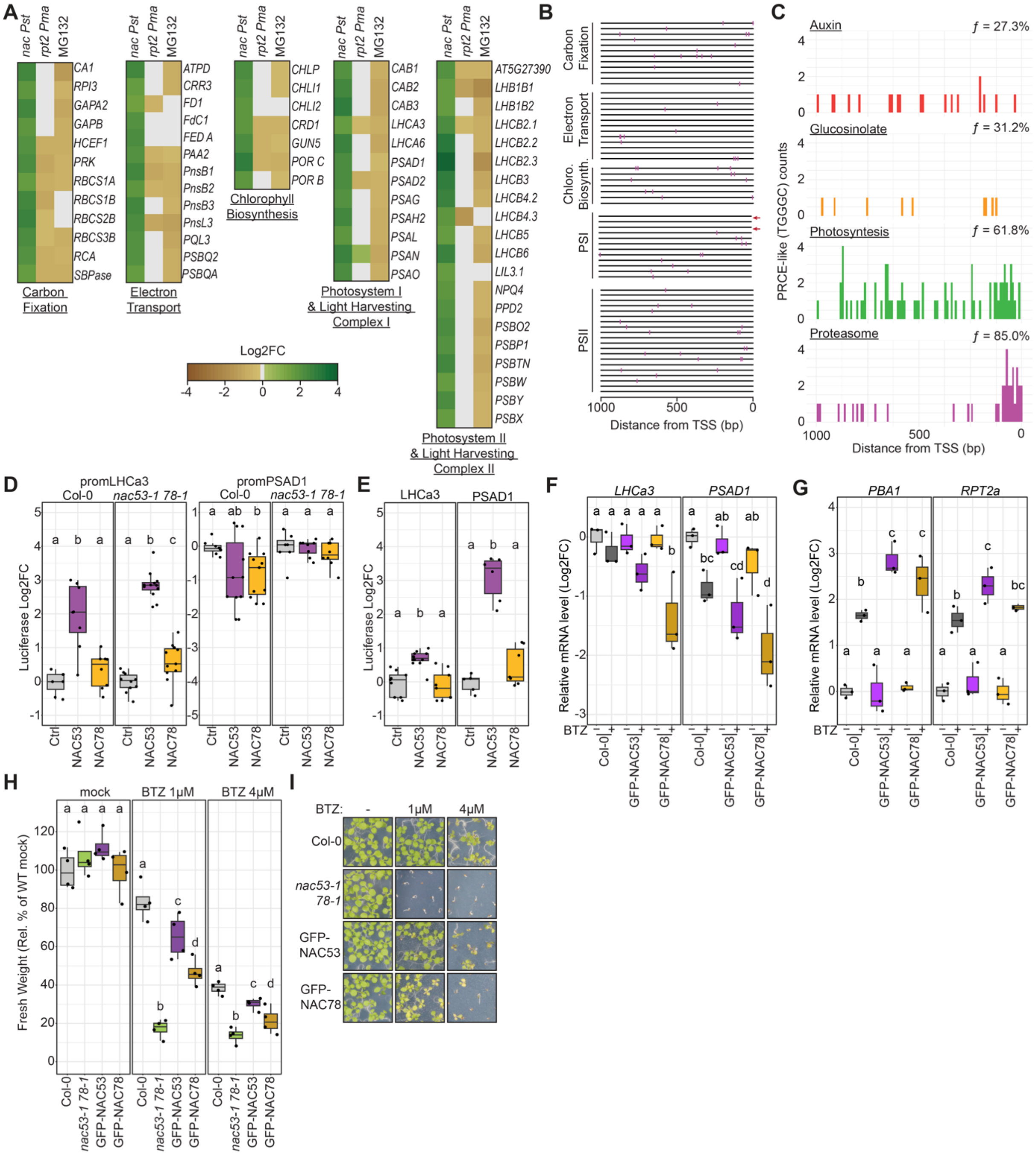


**Extended data Fig.5. Characterization of the NAC53/78-PRCE module in PhANGs promoters.**
(A) mRNA level Log2FC of the PhANGs clusters from Figure 4L in the 3 transcriptomes used. Genes are separated based on their associated process in the photosynthetic pathway.
(B) Mapping of the PRCE-like motifs in PhANGs promoter regions from panel A. Promoter regions are organized according to the order of the genes from panel A. Purple marks represent the position of the motifs based on their distance from the transcription starting site (TSS). The two red arrows highlight LHCa3 and PSAD1 promoter regions.
(C) Histograms representing the number of occurrences of the PRCE-like motifs across the 4 genes cluster from the network Figure 4L. Each bar represents a 20bp region, frequency *f* of promoters bearing at least one motif is indicated as a percentage.
(D) LHCa3/PSAD1 promoter activity measured by luminescence in Col-0 or *nac53-1 78-1* protoplasts extract transiently co-expressing LHCa3/PSAD1 reporter constructs and NAC53/78ΔTM. Log2FC is calculated from the

luminescence level of protoplasts expressing the reporter construct alone. Replicates are a pool of at least two independent experiments. Letters indicate the statistical group assessed by pairwise Wilcoxon-Mann-Whitney test (p value < 0.05).

(E) Promoter activity as calculated in panel D for LHCa3/PSAD1 promoters deleted for their respective DNA regions from Figure 5B. Replicates are a pool of at least two independent experiments. Letters indicate the statistical group assessed by pairwise Wilcoxon-Mann-Whitney test (p value < 0.05).

(F-G) Boxplot representing the Log2FC mRNA level the indicated transcripts in Col-0, GFP-NAC53 or GFP-NAC78 seedlings treated with BTZ 10µM or mock solutions for 6h. Log2FC is calculated from Col-0 mock. Letters indicate the statistical group assessed by pairwise Wilcoxon-Mann-Whitney test (p value < 0.05).

(H) Fresh weight of seedlings from indicated genotypes grown under the indicated treatments at 10 days after germination. Letters indicate the statistical group assessed by pairwise Wilcoxon-Mann-Whitney test (p value < 0.05). Experiment was repeated 3 times with consistent results.

(I) Representative phenotypic pictures related to panel D.

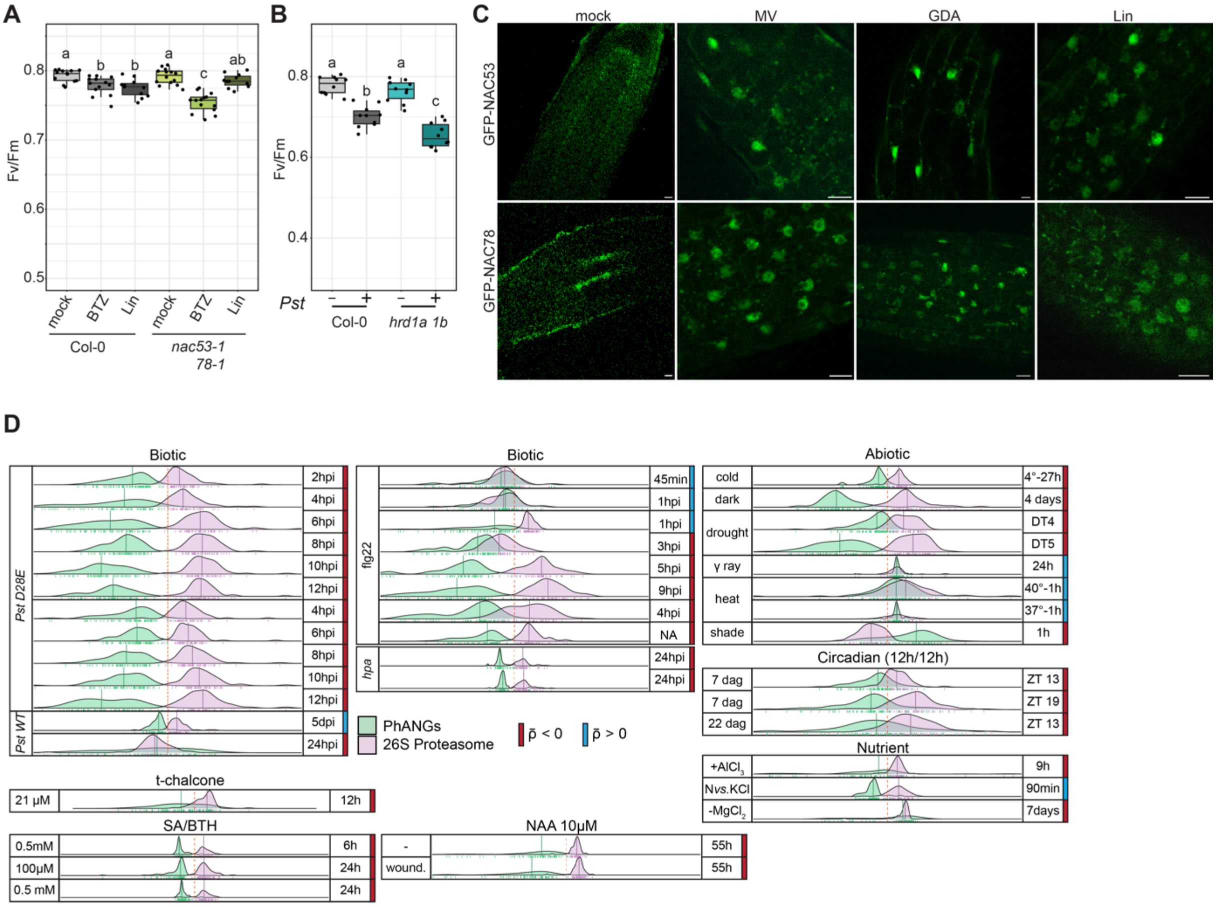

**Extended data Fig.6. The proteasome autoregulatory feedback loop transcriptional signature is a systemic response to environmental cues.**

(A) PSII activity measurement on Col-0 and *nac53-1 78-1* treated as indicated in Figure 6A. Replicates are a pool of 3 independent experiments. Letters indicate the statistical group assessed by pairwise Wilcoxon-Mann-Whitney test (p value < 0.05).

(B) PSII activity measurement on Col-0 and *hrd1a hrd1b* adult plant leaves inoculated with *Pst* or mock solutions for 24h. Letters indicate the statistical group assessed by pairwise Welch t.test (p value < 0.05). The experiment was repeated 2 times with consistent results.

(C) Confocal microscopy pictures of transgenic GFP-NAC53/78 *A. thaliana* roots exposed to mock treatment, MV 100µM, GDA 100µM and Lin 100µM for 2h. The treatments were repeated at least three times with similar observations.

(D) Density functions of the 54 26S proteasome genes and 68 PhANGs transcripts (Log2FC) in the different transcriptome projects. Log2FC limits are -10 to +10. Treatment types and relevant condition information are provided in the figure. Plots are separated by treatment groups and subdivided by experiments.

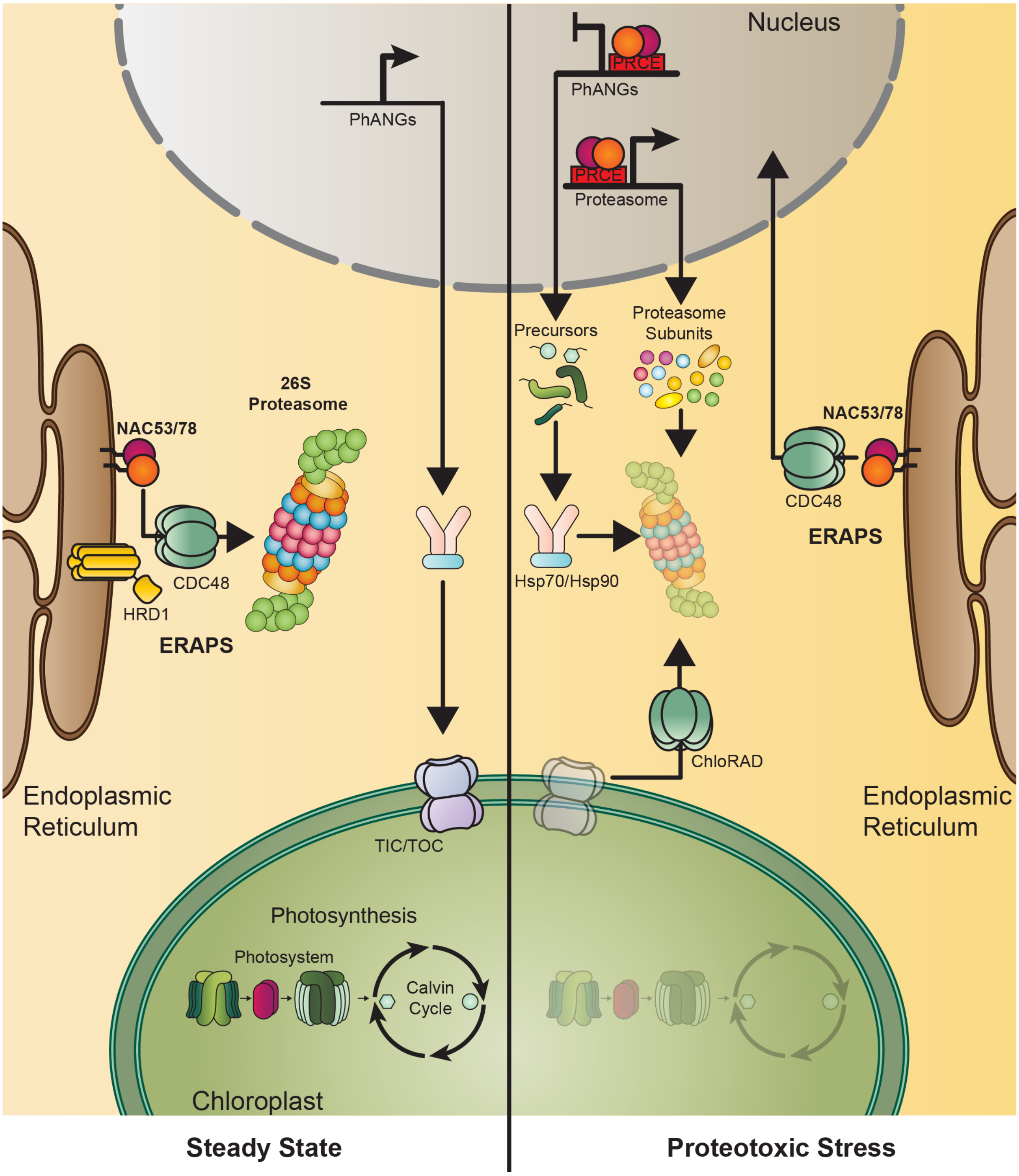


**Extended data Fig. 7. ER-anchored protein sorting (ERAPS) controls the fate of two proteasome activators for intracellular organelle communication during proteotoxic stress.**
In steady state conditions, the ER-anchored protein sorting system (ERAPS) promotes the constitutive degradation of NAC53/78 via the 26S proteasome. Meanwhile, PhANGs expression is activated and subsequent chloroplastic import permits the maintenance of active photosynthesis. Upon proteotoxic stress, the chloroplast import machinery is targeted for proteasomal degradation. This leads to an accumulation of PhANGs precursors which are therefore subjected to proteasomal degradation, inducing proteotoxicity. Thus, to avoid proteotoxicity, the ERAPS system facilitates the nuclear translocation of NAC53/78, to activate the production of a new proteasome complex and to repress PhANGs expression to mitigate substrate accumulation.
